# A guide to sampling design for GPS-based studies of animal societies

**DOI:** 10.1101/2022.01.29.478280

**Authors:** Peng He, James A. Klarevas-Irby, Danai Papageorgiou, Charlotte Christensen, Eli D. Strauss, Damien R. Farine

**Affiliations:** Department of Collective Behaviour, Max Planck Institute of Animal Behavior, Konstanz, Germany; Centre for the Advanced Study of Collective Behaviour, University of Konstanz, Konstanz, Germany; Department of Biology, University of Konstanz, Konstanz, Germany; Department of Evolutionary Biology and Environmental Science, University of Zurich, Zurich, Switzerland; Department of Migration, Max Planck Institute of Animal Behavior, Radolfzell, Germany; Division of Ecology and Evolution, Research School of Biology, Australian National University, 46 Sullivans Creek Road, Canberra, ACT 2600, Australia; Department of Ornithology, National Museums of Kenya, P.O. Box 40658-001000, Nairobi, Kenya; Mpala Research Centre, Nanyuki, 10400, Kenya

**Keywords:** Animal societies, Bio-logging, GPS sampling, GPS tracking, Group and collective behaviours, Group size, Social interactions, Social structure

## Abstract

GPS-based tracking is widely used for studying wild social animals. Much like traditional observational methods, using GPS devices requires making a number of decisions about sampling that can affect the robustness of a study’s conclusions. For example, sampling fewer individuals per group across more distinct social groups may not be sufficient to infer group- or subgroup-level behaviours, while sampling more individuals per group across fewer groups limits the ability to draw conclusions about populations. Here, we provide quantitative recommendations when designing GPS-based tracking studies of animal societies. We focus on the trade-offs between three fundamental axes of sampling effort: 1) sampling coverage—the number and allocation of GPS devices among individuals in one or more social groups; 2) sampling duration—the total amount of time over which devices collect data; 3) sampling frequency—the temporal resolution at which GPS devices record data. We first test GPS tags under field conditions to quantify how these aspects of sampling design can affect both GPS accuracy (error in absolute positional estimates) and GPS precision (error in the estimate relative position of two individuals), demonstrating that GPS error can have profound effects when inferring distances between individuals. We then use data from whole-group tracked vulturine guineafowl (*Acryllium vulturinum*) to demonstrate how the trade-off between sampling frequency and sampling duration can impact inferences of social interactions and to quantify how sampling coverage can affect common measures of social behaviour in animal groups, identifying which types of measures are more or less robust to lower coverage of individuals. Finally, we use data-informed simulations to extend insights across groups of different sizes and cohesiveness. Based on our results, we are able to offer a range of recommendations on GPS sampling strategies to address research questions across social organizational scales and social systems—from group movement to social network structure and collective decision-making. Our study provides practical advice for empiricists to navigate their decision-making processes when designing GPS-based field studies of animal social behaviours, and highlights the importance of identifying the optimal deployment decisions for drawing informative and robust conclusions.

## 1. INTRODUCTION

Studying animal societies requires collecting both individual-level (what is the animal doing?) and relational (what is the relationship between animals?) data (Altmann 1974; Krause *et al*. 2013). Prior to data collection, researchers have to make decisions about how to employ finite resources to adequately sample social behaviours. While sampling design is a major consideration across all fields of biology, it is critical when studying social behaviour because the number of possible inter-individual relationships increases exponentially with the study population size (Davis, Crofoot & Farine 2018). Further, social relations can change over time (Pinter-Wollman *et al*. 2014; Gil *et al*. 2018), and vary across social organizational scales and social systems. Thus, studying social behaviour benefits from sampling individuals simultaneously, frequently, broadly, and over large temporal scales. Given their ability to collect large amounts of data simultaneously, automated sampling methods are increasingly used to remotely collect data on animals (Krause *et al*. 2013; Nomano *et al*. 2014; Wilmers *et al*. 2015; Ferreira *et al*. 2020; Gupte *et al*. 2021; Keitt & Abelson 2021; Shizuka *et al*. 2021; Smith & Pinter-Wollman 2021). The proliferation of these novel sampling techniques motivates a need for practical guidance to assist empiricists in designing behavioural sampling regimes that are best suited for delivering new insight about animal societies.

Remote tracking of animals dates back to the 1960’s (Craighead 1982). However, the large-scale uptake of GPS for animal tracking has come in the past two decades, following major breakthroughs in GPS-based technologies, including improved precision (Frair *et al*. 2010; Dujon, Lindstrom & Hays 2014), lower costs (Rodgers 2001), and greater battery efficiency and weight reduction (Kays *et al*. 2015). Such widespread applications of animal GPS tracking have also stimulated advances in analytical tools (Robitaille, Webber & Vander Wal 2019; Joo *et al*. 2020; Silva *et al*. 2022), making it increasingly possible to not only study where individuals distribute themselves in space (e.g., for wildlife management or monitoring purposes), but also how animals move with respect to each other at various scales (Robitaille *et al*. 2021). For example, studies have used GPS tracking data to infer the likelihood of encounters between conspecifics (Noonan *et al*. 2021) and the ways in which animals influence each other’s movements (King *et al*. 2012; Long *et al*. 2014; Schlägel *et al*. 2019; Milner, Blackwell & Niu 2021). Such studies have demonstrated the potential for using GPS tracking to study animal social behaviours (Juang *et al*. 2002; Hebblewhite & Haydon 2010; Spiegel *et al*. 2016; Sih *et al*. 2018; Spiegel *et al*. 2018; Gilbertson, White & Craft 2021), and the interface of social behaviour with movement ecology (Torney *et al*. 2018; Westley *et al*. 2018; He, Maldonado-Chaparro & Farine 2019).

GPS telemetry explicitly captures when and where individual animals go, opening new avenues of research into how animals interact with their social and ecological environment (Cagnacci *et al*. 2010; Tomkiewicz *et al*. 2010; Kays *et al*. 2015; Leu *et al*. 2016; Webber *et al*. 2022). GPS devices can substantially increase the volume and dimension of data collected relative to human observers (Crofoot 2021). Importantly, GPS devices can be deployed on more individuals than could possibly be observed simultaneously, potentially capturing fine-scale social processes such as group movement and collective decision-making (Strandburg-Peshkin *et al*. 2015). Studies over the last decade have established simultaneous GPS-based tracking as a productive approach for generating new insights into animal social ecology (Dell’Ariccia *et al*. 2008; Nagy *et al*. 2010; Lührs & Kappeler 2013; Strandburg-Peshkin *et al*. 2015; Leu *et al*. 2016; Spiegel *et al*. 2016; Springer *et al*. 2016; Farine *et al*. 2017; Papageorgiou *et al*. 2019; Wielgus *et al*. 2021). Yet, despite its strengths, GPS tracking is far from being a panacea for challenges related to behavioural data collection (Hebblewhite & Haydon 2010). The cost of GPS devices (and data transfer), the challenges of deployment, and technological limitations can all constrain study design.

To effectively apply GPS tracking to study social behaviours, researchers need to tailor sampling design to research questions and social systems. Animal social behaviours span various spatial and temporal scales (Robitaille *et al*. 2021) and levels of social organization (Wilson 1975; Aureli *et al*. 2008; Prox & Farine 2020), such as how one individual interacts with another, how individuals form groups, how groups themselves interact, how populations are socially structured and how these dynamics change over time (Lukas & Clutton-Brock 2018; He, Maldonado-Chaparro & Farine 2019; Kappeler 2019). Optimal GPS study design strategies will therefore depend on the question being investigated—for example, tracking all individuals in one social group is useful for sampling interactions among group-mates but not among groups. Another important consideration is that some animals associate almost exclusively with the same set of individuals (e.g., within territory boundaries, Bishop & Groves 1991; Jordan, Cherry & Manser 2007), whereas others form open societies where group membership and size change frequently (Aureli *et al*. 2008; Connor & Krutzen 2015; Farine *et al*. 2015). By influencing how often individuals are found together, these dynamics shape how GPS tracking can be employed to study social behaviours.

In this paper, we first summarize the common trade-offs faced by researchers using GPS tracking to study animal societies, focusing on three key axes: sampling coverage, sampling duration, and sampling frequency. We then use field tests of GPS tags to demonstrate how sampling regimes can affect error, and subsequently our ability to infer social behaviours. Finally, we use data from two whole-group GPS tracking datasets of vulturine guineafowl (*Acryllium vulturinum*), together with data-informed simulations, to investigate how sampling coverage affects inference of social interactions and social metrics within different animal social groups. In doing so, we provide practical advice for the design and implementation of GPS-based studies of social behaviours (Table 1).

**Table 1:**
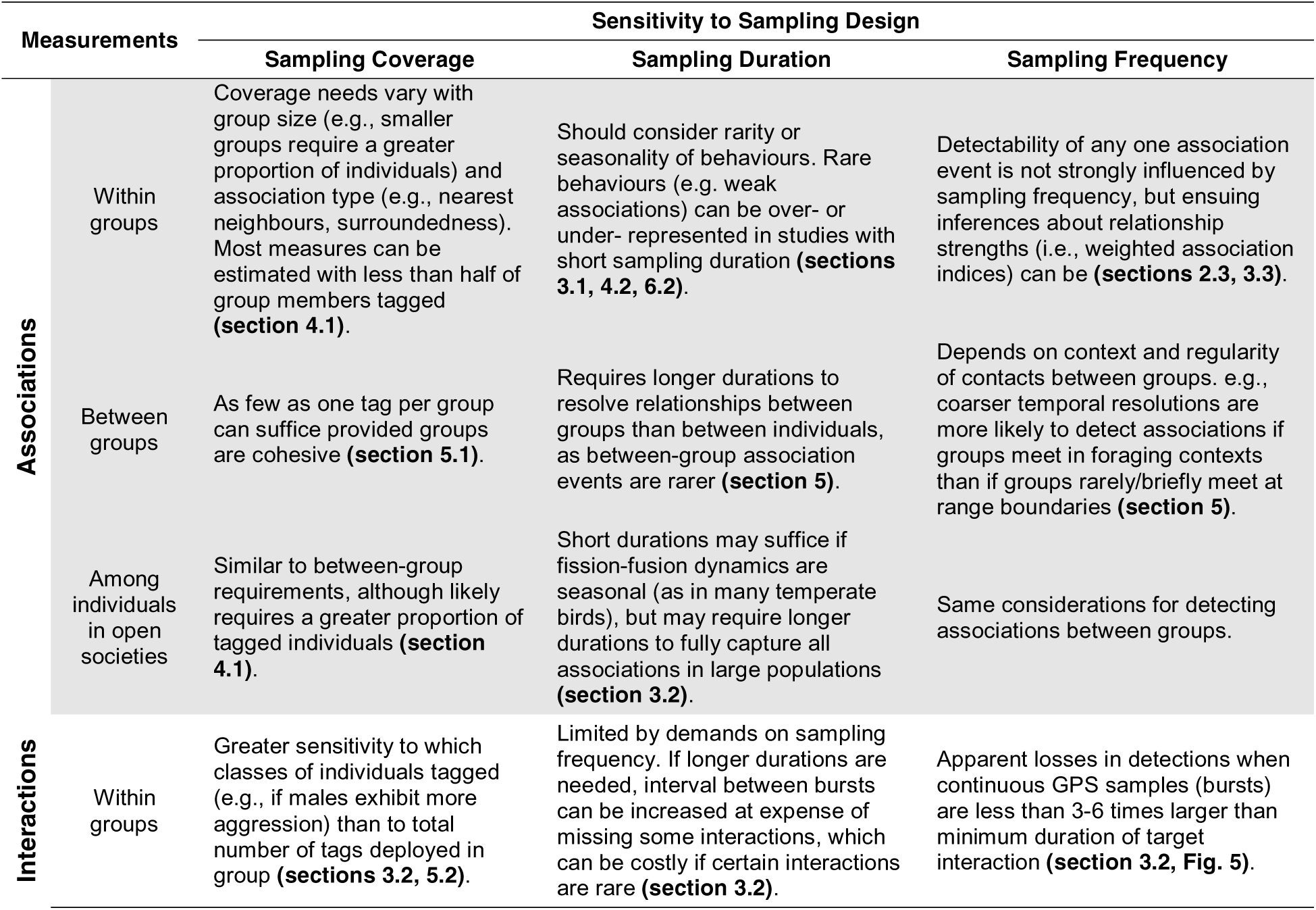
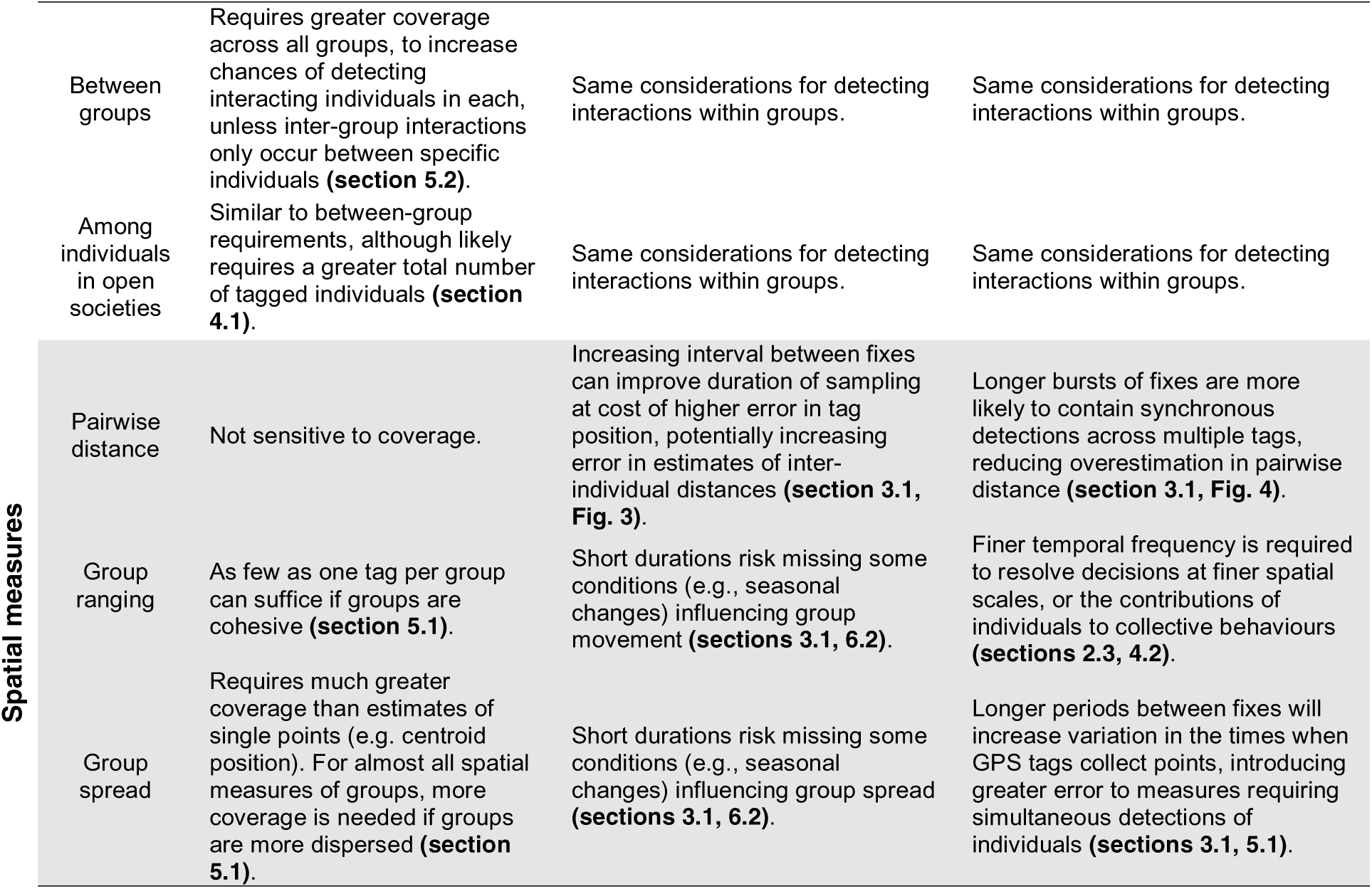
Summary recommendations for sampling coverage, duration, and frequency when measuring associations, interactions, and space use across different levels of social organisation.

## 2. KEY TRADE-OFFS IN GPS-BASED STUDIES OF SOCIAL BEHAVIOURS

Many factors should be considered when designing GPS-based tracking studies of social behaviours (Fig. S1). Among these are ethical and practical limitations when capturing and tagging animals, and ongoing research and development is aimed at understanding and minimizing these limitations (Portugal & White 2018; Klegarth *et al*. 2021; Portugal & White 2021; Wilson *et al*. 2021). Here, we assume that researchers follow ethical and legal guidelines for deploying GPS devices and focus on providing guidance on how to design the deployment of GPS devices to optimize data collection for addressing different research questions (Fig. 1).

**Figure 1.**
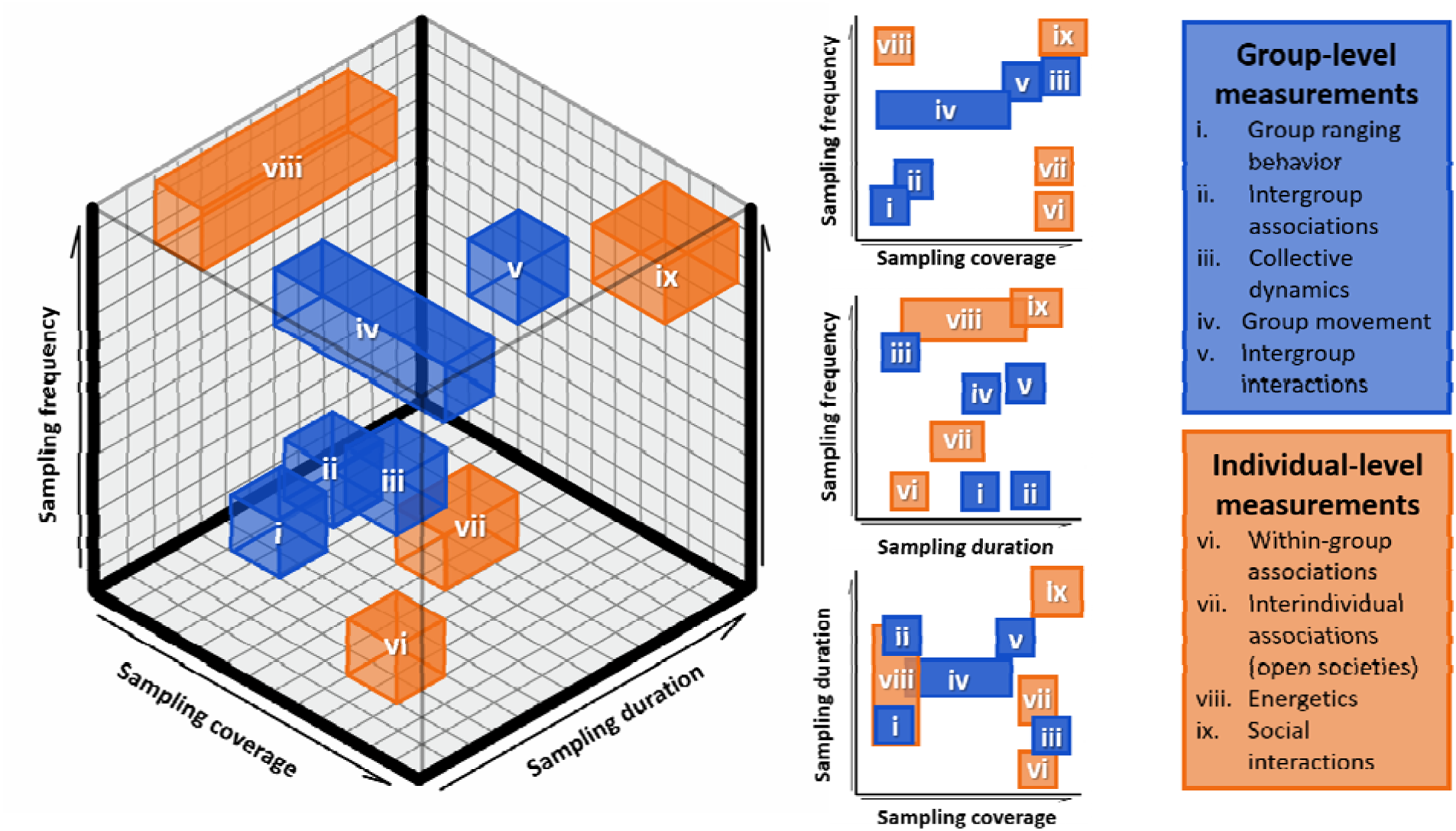
Trade-offs across three fundamental dimensions of GPS sampling design when studying animal societies. The types of questions that can be answered using GPS tracking are broadly governed by three key axes of study design: sampling coverage—the number and distribution of GPS device throughout a study population, sampling duration—the total amount of time over which devices collect data, and sampling frequency—the temporal resolution at which devices collect data. Social behaviour can typically be thought of as playing out at group (blue boxes) and individual (orange boxes) levels. Her we illustrate how, within each level, different measurements (i-ix) require differing levels of investment in each axis, and often face trade-offs across axes to capture specific behaviours of interest. For example, group ranging behaviour (how large is a home range) requires lower sampling frequency or coverage than group movement (how fast do groups move) or collective dynamics (how do groups make decisions). Further, rarer social behaviours (e.g. inter-group associations) require longer durations than more frequent social behaviours (e.g. within-group associations).

### 2.1 Sampling coverage

All GPS-based studies face financial and practical constraints on the number of devices that can be deployed. Having more devices (relative to the study population size) will naturally provide more data for more robust inferences of social behaviours. However, deploying more devices is not only financially costlier, it can also introduce a number of logistical challenges in terms of data-recovery (see 2.4). Therefore, researchers need to strategically decide how many devices to deploy, and how broadly to distribute them (i.e., ‘sampling coverage’).

Sampling coverage requirements will depend on the scale of social organization in relation to research questions and whether a study focuses on social structure (e.g., patterns of association or interactions) or processes (e.g., transmission or collective behaviour). Structure-focused questions, such as whether individuals have preferred or avoided social relationships (e.g., Cantor *et al*. 2012; Best *et al*. 2014; Gero, Gordon & Whitehead 2015), require broad, but not necessarily exhaustive, deployments of devices—it is fine to not capture every single pair of associates, as long as a range of association strengths are represented. The number of devices to be deployed will of course also depend on social scale (e.g., group vs population). In terms of allocation, randomly deploying GPS devices to individuals should be suitable for testing for the presence of preferred social relationships. Questions addressing larger population-scale structure, such as whether there are distinct social communities within a population (e.g., Shizuka & Farine 2016), may require distributing GPS devices more broadly to distinguish higher levels of social organisation, while those seeking finer-scale structure, such as who are an individual’s closest associates, may require more focused deployments.

Process-focused questions require thorough coverage of social relationships, which can be very device-intensive. Without a relatively complete representation of social connections, inferences may fail (Perkins *et al*. 2009; Wild & Hoppitt 2019). For example, observing a pathogen spreading from individual A to B to C, with edges A–B and B–C, would fail to reveal socially-mediated transmission if A and C become infected and B is missing from the study. As such, process-focused questions may need to be studied at smaller scales, where higher proportions of individuals can be tracked, or to focus on interactions among larger social organizational units (e.g., where an entire group can be represented by one device).

### 2.2 Sampling duration

Sampling duration is fundamentally limited by the battery life of GPS devices (Fig. S1). Power is consumed each time a device switches on to collect an animal’s position, leading to a major trade-off between the sampling frequency and the longevity of battery life that determines sampling duration. For example, sampling of GPS collars fitted to wild olive baboons (*Papio anubis*) lasted 15 to 30 days at 1 Hz (Farine *et al*. 2017), whereas those fitted to elephants (*Loxodanta africana*) operate at much lower sampling frequencies but can last many years (Hahn *et al*. 2022; also because larger body sizes allow for larger batteries). Some GPS devices can partly overcome this challenge by using solar panels to recharge batteries, by applying programming settings such as using geofencing to spatially vary sampling regimes (Sheppard *et al*. 2015), or by using acceleration sensors to switch off/on GPS devices when animals are (in)active (Brown *et al*. 2012). Limitations to battery (and sometimes on-board data storage) capacity means that researchers often need to trade-off between sampling more intensively (higher sampling frequency) versus sampling for longer (see the duration-frequency trade-off below). The key consideration is therefore whether a given sampling duration can reveal fine-scale social processes and the long-term factors that govern them.

### 2.3 Sampling frequency

Sampling frequency determines the temporal resolution at which behavioural inferences can be made. Animal movement is inferred from temporally sequential fixes, so the more fixes are collected within a given time window, the more complete the information is about where the individual has been and, potentially, what it has been doing (Wang 2019). In most vertebrates, sampling frequency at 1 Hz (1 fix/s) captures rich information about where individuals have moved and how (Ryan *et al*. 2004). However, such high sampling frequencies are energetically more demanding, generally at the cost of sampling duration. To save battery life, studies often opt for lower sampling frequencies, such as 10 fixes every 5 minutes (e.g., vulturine guineafowl, Papageorgiou *et al*. 2019), 1 fix every 30 minutes (e.g., swamp wallabies *Wallabia bicolor*, Fischer *et al*. 2018), or 1 fix every hour or more (e.g., African elephants, Wittemyer *et al*. 2008). However, while allowing behavioural characterizations over larger temporal scales (e.g., over years), lower sampling frequencies may not adequately capture social behaviours that are realized at finer temporal scales (Haddadi *et al*. 2011; Wilson *et al*. 2014), such as how collective movement decisions are made. From a social network perspective, the frequency-duration trade-off will depend on whether the aims are to collect social interactions or associations.

The restrictions of lower sampling frequency can be partly addressed by inferring intermediate locations (i.e., an animal’s position in the time between fixes) using approaches such as spatial interpolation (Strandburg-Peshkin *et al*. 2015; Hirakawa *et al*. 2018), smoothing spline models (Whetten 2021), or dead reckoning (Dewhirst *et al*. 2016). Related to this, it should also be noted that battery consumption does not scale linearly with increasing sampling frequency. GPS devices use energy each time they (re)boot and search for satellites, which typically takes longer with sparser sampling regimes or under when animals are under canopy cover, meaning that the same battery charge collects fewer data points at longer sampling intervals. Device may also not switch off during more intensive regimes. For example, 0.2–0.5 Hz may use as much battery as 1 Hz (but 1 Hz may have greater battery costs for data transfer).

### 2.4 Further practical considerations

It should be noted that other practical issues can shape decisions on GPS sampling design (Fig. S1). Among these, GPS accuracy (i.e., the extent to which the fixes of a GPS device capture the actual locations on the planet, such as those defined in WGS-84) and/or precision (i.e., the extent to which fixes of a GPS device consistently agree with each other; see Yoshimura & Hasegawa 2003) pose a fundamental challenge for inferring animal behaviours (Frair *et al*. 2010; Adams *et al*. 2013; Clements *et al*. 2022).

Apart from GPS accuracy/precision, researchers also need to consider data retrieval, which can become cumbersome when deploying many devices simultaneously. Deploying more devices will increase costs if data are retrieved via mobile phone networks, download time if data are retrieved via radio download, or re-trapping/relocation effort if data is retrieved by recovering devices. Device data storage capacity may require frequent data retrieval and missed data downloads can lead to data loss (e.g., Strandburg-Peshkin *et al*. 2015 retrieved GPS data everyday to avoid data loss; see also Box 1). Finally, if tags cannot be downloaded often, then they may have insufficient battery life to download very large amounts of data.

## 3. SOCIAL NETWORK STRUCTURES

Animal social networks are often assembled from social associations or interactions (Whitehead 2008; Farine 2015; Farine & Whitehead 2015; Krause *et al*. 2015), both of which can be inferred from GPS data (Robitaille, Webber & Vander Wal 2019; Robitaille *et al*. 2021). In GPS tracking studies, social associations are defined by co-location of individuals, while interactions can be inferred when individuals have a detectable impact on the behaviour of others (e.g., individuals’ movement responses to each other; Long *et al*. 2014; Schlägel *et al*. 2019; Milner, Blackwell & Niu 2021). Several studies have suggested that social networks are quite robust to sampling subsets of the population— Silk *et al*. (2015) estimated that studies sampling 30% or more of a population can generate reliable estimates of individuals’ social network positions—so GPS devices can theoretically be deployed to characterize even large social networks. Subsequent studies highlighted that repeated sampling and avoiding misidentifications are critical for producing accurate networks (Davis, Crofoot & Farine 2018; Sunga, Webber & Broders 2021)—both are the strengths of GPS tracking data. However, GPS positions are also prone to error and sampling discontinuity (i.e. due to the frequency-duration trade-off). Below we discuss sources of GPS error and some implications of sampling design when inferring associations and interactions.

### 3.1 Effects of sampling design on GPS accuracy, precision, and data synchronisation

First, it is important to understand sources of positional error (GPS accuracy) when taking a GPS fix. To test how habitat affects error, we set out ten e-obs 15g solar GPS tags (five in open habitat and five in closed canopy conditions), all programmed to record 10 fixes (at 1Hz)) every 5 minutes from 6am to 7pm. We used the median position for each tag (after removing outliers) to estimate the true position, and calculated the latitudinal and longitudinal distance for each fix from this true position as the error. Outliers (see Gupte *et al*. 2022 for a guide on how to deal with outliers) were defined as points beyond two times the 99.9^th^ percentile of distances for the open area tags (36.68 m). We found that the distribution of positional errors is generally leptokurtic (more peaked than a Gaussian distribution), with the distribution of errors being ‘broader-shouldered’ (i.e. less leptokurtic) under canopy cover (Fig. 2). As with previous work (Langley 1996; Frair et al. 2010; Crofoot 2021), we also found that canopy cover delayed acquisition of satellites, even with a 5-minute inter-burst interval. This can have important consequences when collecting social data, as it can decrease the temporal synchronisation data collected across individuals (Fig. 2c).

**Figure 2.**
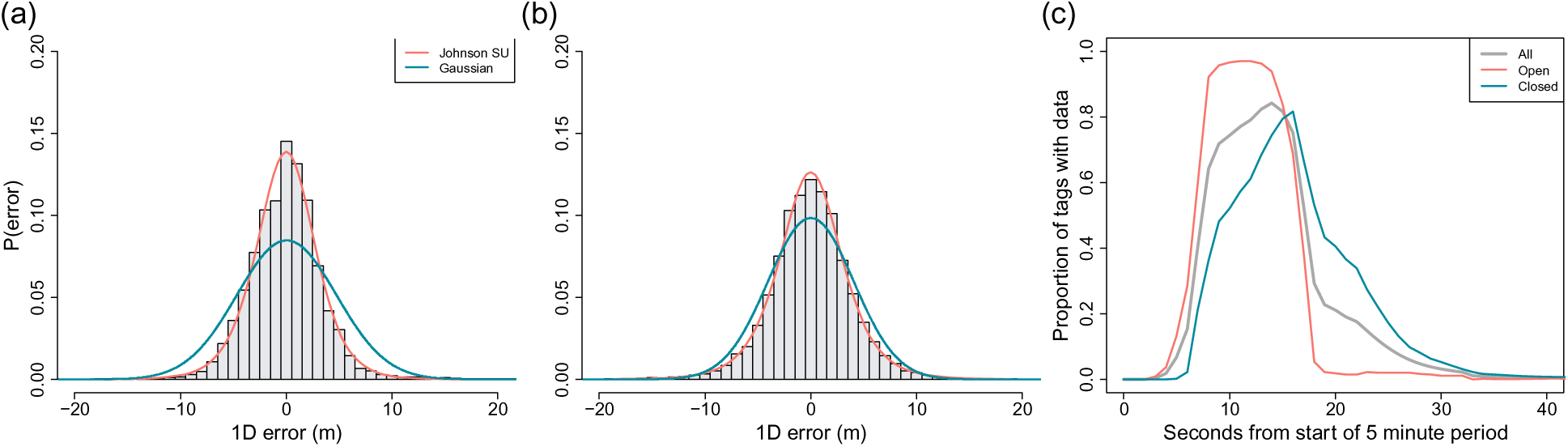
GPS accuracy and synchronisation vary with habitat. Distributions of error estimated from five GPS-tags placed in (a) open vs. (b) closed canopy conditions, collecting a burst of 10 points every 5 minutes. Both probability densities follow a leptokurtic distribution, but the tighter shoulders around the open condition suggests that fewer large errors occur compared to closed canopy. Red line shows the fit of a Johnson SU-distribution. Blue lines show the best fitting normal distribution. (c) Synchronisation in data collection for tags with 5-minute inter-burst intervals, showing a delay (and reduced synchronisation) under closed canopy cover.

In theory, one way to reduce GPS error is to use continuous sampling. This is because devices can reuse satellite information from previous fixes (e-obs GmbH, pers. comm.). In contrast, switching off devices between consecutive fixes will generally result in lower accuracy, with error increasing as the between-sample gap increases. To verify this, we programmed four 15g e-obs solar tags to collect burst of 10 consecutive fixes at different inter-burst intervals (5, 10, 30 and 60 min). We placed these in the same position for 8 consecutive days, collecting data from 6am to 7pm each day. From these data, we calculated the median position (estimated true location) for each tag, and estimated the error around this location for each fix. We found that the mean error around this point (Fig. 3a) and the percentage of outliers (Fig. 3b) decreased with an increasing number of fixes and with shorter inter-burst intervals. We also show that the further an animal moves between each burst (simulated by displacing the tags 10 or 60 meters between each burst), the larger the error (Fig. 3a). Such checks are worth conducting when designing a sampling regime as positional accuracy will vary across different tag types, conditions, or behaviours (Gunner *et al*. 2022). We also found that error information derived from the tag itself may not be informative (see Supplementary Material section 2).

**Figure 3.**
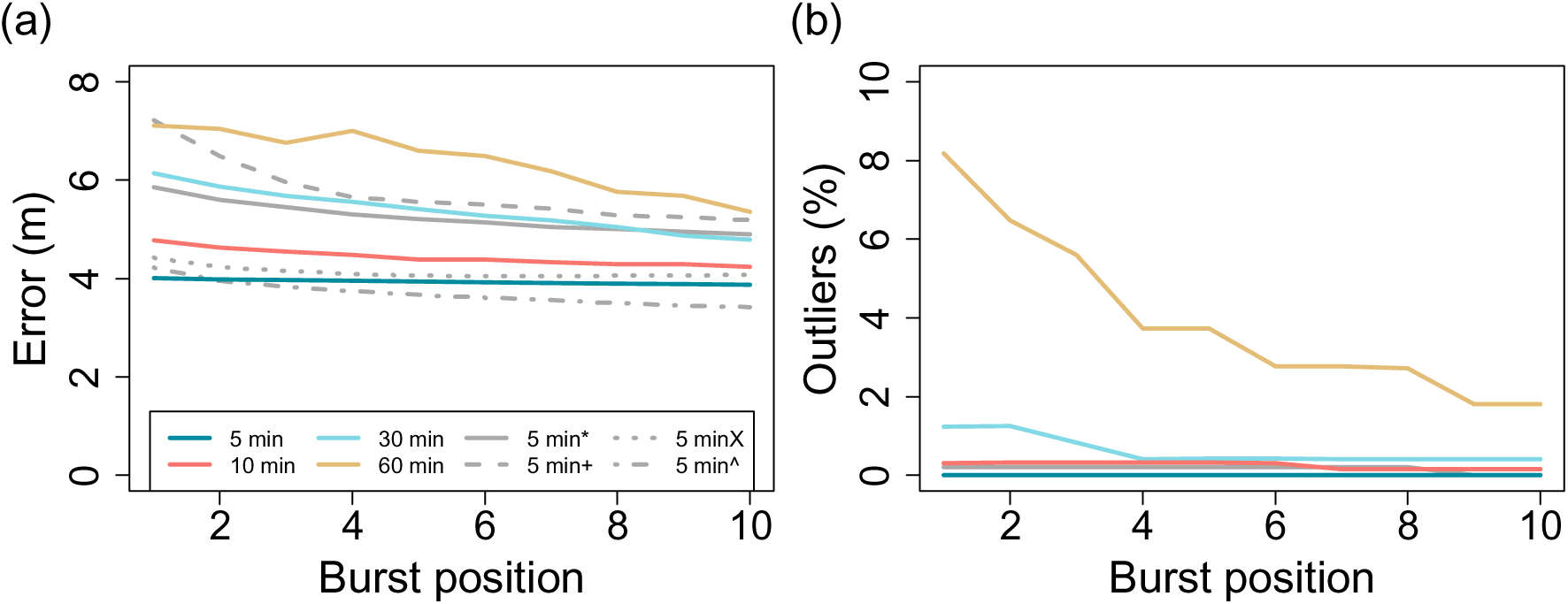
GPS accuracy under different sampling regimes and conditions. (a) Mean error around ‘true location’ and (b) percentage of outliers decrease with shorter inter-burst intervals (colours) and burst length (1st to 10th fix). 5 min* refers to the 5-minute inter-burst interval under canopy cover, 5 minX refer to tags that moved 10 m between bursts (open habitat), 5 min+ refer to tags that moved 60 m between samples (open habitat), and 5 min^ refer to tags in stationary positions 50cm off the ground (open habitat).

### 3.2 Estimating social associations

Associations can be defined by applying spatiotemporal co-location rules to individuals’ GPS fixes. A common approach—at least for terrestrial animals—is to set a horizontal distance threshold under which individuals are considered associated. While thresholds should consider the biology of the study organisms, the ability to apply meaningful thresholds will be impacted by GPS precision. Given a satisfactory threshold, the decision is then whether to sample more intensively (higher sampling frequency) or to sample for longer (longer duration).

Defining a biological threshold for associations can be based on traditional observation of the study organism, through data-driven studies of these thresholds (Haddadi *et al*. 2011), or by using statistical approaches (Papageorgiou *et al*. 2019). An example of the latter is to interrogate the distribution of inter-individual distances and extract the first mode, which likely represents individuals that are socially associating. In larger groups, associations can be defined following a chain rule, where all individuals that form part of a connected component are considered as being associated (e.g., if A–B and B–C, then A–C even if A and C are beyond the threshold). There are also tools (Ansari *et al*. 2020) for identifying clusters of GPS fixes to infer behavioural events or states (e.g., kills by predators or behavioural activity states). However, few have been extended to detect clustered social associations (i.e., inferring the presence of social aggregations) across time (but see Kalnis, Mamoulis & Bakiras 2005).

The chosen threshold then needs to take into consideration GPS accuracy (positional error), because positional error leads to overestimates of inter-individual distances (see Supplementary Materials section 3), i.e. reduced precision. This is due to the circular nature of the GPS error. Imagine two individuals resting at a set distance apart. If the positional error is distributed around these individuals in a circular fashion, then the proportion of the areas in these circles in which a simultaneous detection of both individuals is at, or below, their actual distance is smaller than the area in which the detections will be above their actual distance (when two individuals are at exactly the same position, the distance can only be greater or equal to 0, and never less). Supplemental materials section 3 demonstrates this and shows, using simple simulations, that the extent of the overestimate of between-device distance relative to true distance increases with positional error and decreases with true distance.

To verify the predicted relationship between true distance and distance overestimation with tag error, we placed five 15g e-obs solar tags in a straight line at set spacings (0, 1, 3, 6 and 9 meters; resulting in a matrix of 10 pair-wise distances between 1 and 9 meters), recording a burst of 10 fixes (at 1Hz) every 5 minutes. We also used the data from the tags that moved between two fixed positions (10 m apart and 60 m apart) between every burst, as well as pairs of tags attached at a fixed 1 meter distance while being moved either side-by-side or front-to-back. We then calculated the distance between time-matched GPS fixes for each pair of tags and compared this with their true distance. Further, we calculated the theoretical overestimate for each underlying true distance, by sampling pairs of points from the observed error distribution from each device. Our data confirmed that between-device distances are generally overestimated, and more so at shorter true distances (Fig. 4). However, this relationship is dampened relative to the theoretical prediction because tag precision is greater than tag accuracy, which is because the direction of the error in simultaneous GPS fixes are correlated, with closer devices having more correlated errors (Fig. 4). This is likely because tags that are closer together share the same satellites to obtain their position, resulting in the direction of their error being the same. We confirmed this spatiotemporal autocorrelation in GPS error by randomising the matched GPS-fixes (to break the correlations), with overestimation error matching the theoretical prediction more closely (see Supplementary Material section 4). This proximity bias has implications for inferring social associations or interactions defined by inter-individual distance using GPS data, which will depend on the size of the animal or the nature of the social association (e.g., co-foraging vs. grooming). For example, studies may need to set their distance threshold higher than the biologically meaningful threshold if they wish to capture true association rates (see Supplementary Material section 5). This effect will be more pronounced in smaller species (or for smaller biological thresholds) and for species (or activities) that are affected more by cover (e.g., grooming under tree canopies; Fig. 4C).

**Figure 4.**
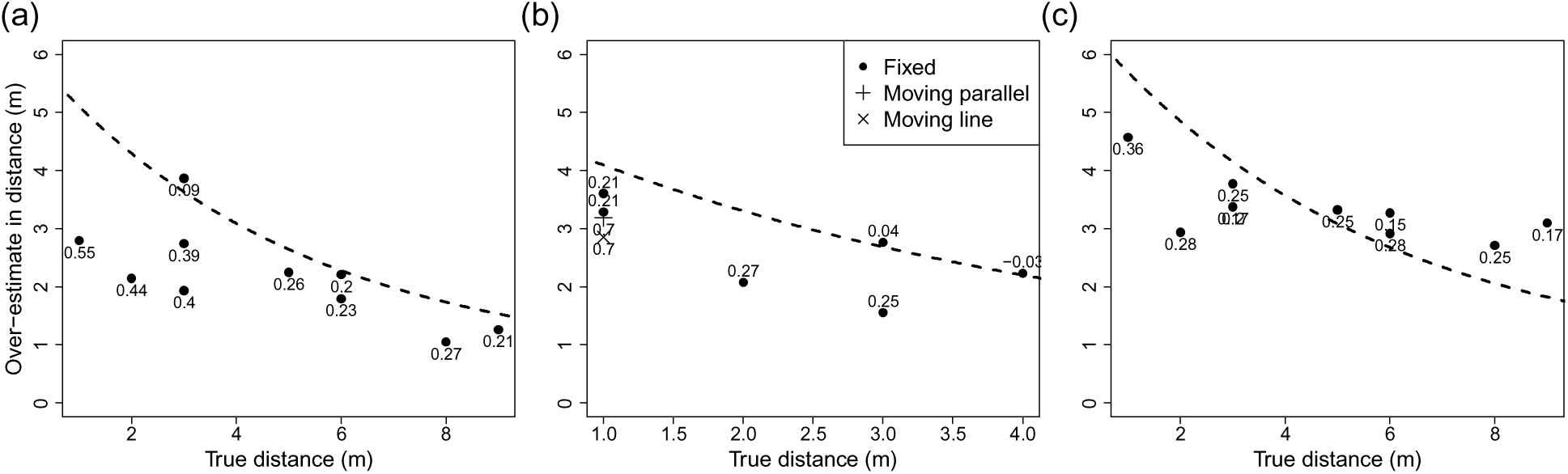
The overestimation in inter-individual distance as a function of the true distance and habitat cover. (a) GPS-based distances consistently overestimate the true tag distance, but do not do so as much as predicted in theory (dashed line, which was calculated by sampling points with a given true distance and an error drawn from the empirical tag error distributions; see Fig. 2). This is likely because the pairwise correlations (numbers in plot) in errors between tags (when using simultaneous data points) decreases as a function of between-tag distance. Breaking this spatiotemporal correlation confirms the theoretical predicted relationship (Fig. S3). (b) Tags that are above ground and tags being ‘on the move’ marginally reduces GPS error. (c) Canopy cover increases error, and also causes overestimations to deviate from the predicted relationship at larger true distances.

Studying social associations also requires considering sampling frequency. The choice of sampling frequency will need to consider the biological context being studied, especially how dynamic the social structure is (Hobson, Avery & Wright 2013; Farine 2018) and when it changes. Imagine that a GPS device has the capacity to collect 4380 fixes. We could opt to collect one fix per hour over 12 months (for 12 hours a day), or to collect one fix every 5 minutes for one month (i.e., two extreme ends of the duration/frequency trade-off). If we are studying animals in a seasonal system, the 12 months data would be able to capture the general social environment that each individual has experienced over the course of the year, but might frequently miss rarer behaviours or associations. By contrast, a one-month deployment could only capture the social environment in that specific month, meaning that many social connections that formed under different conditions could not be observed. A one-month deployment would also likely underestimate the distinct number of connections in an individual’s social environment, thereby overestimating the importance of those that were detected.

### 3.2 Inferring social interactions

An emerging application of GPS tracking is to infer social interactions among individuals. Unlike associations, which are readily characterised at low sampling frequencies from simultaneous detections at single time points, interactions typically involve a time component that requires sequential detections collected simultaneously across individuals. For example, two individuals detected next to one-another may be grooming or walking in opposite directions. Dominance interactions can be thus be inferred from approach-avoid interactions (A approaches B and B moves away) and chases (Strandburg-Peshkin *et al*. 2015). Thus, inferring interactions will require not only considering GPS accuracy and precision, but also careful consideration of the sampling frequency-duration trade-off.

The ability to detect interactions from GPS tracking data will first depend on the spatial scale over which they occur relative to GPS error. For example, a relative positioning GPS error (i.e., precision) within 1 m allows for detections of displacements that occur over tens of meters in baboons (e.g., Strandburg-Peshkin *et al*. 2015), but make it more challenging to detect those that take place at much smaller spatial scales, as in the vulturine guineafowl (Dehnen *et al*. 2022). One potential solution is to supplement GPS data with accelerometer data to better estimate whether two individuals are actually stationary or not (see section 6.5). In cases where error is unavoidable, GPS tracking is perhaps best-suited to detecting extreme interactions (e.g., chases), as GPS precision is higher when tags are moving (Fig. 4-b; see also in Gunner *et al*. 2022). Such issues aside, detecting interactions is achievable, but we encourage manual verification of inferred interactions by traditional observations (e.g., by recording individual behaviours while GPS tracking). Further, it should be feasible to extend machine learning approaches (Valletta *et al*. 2017; Wang 2019) for detecting patterns in temporal data to extract interactions from GPS trajectories.

Given the need to collect periods of sequential data (i.e., bursts) that can capture the temporal patterns of social interactions, studies aiming to detect interactions face difficult decisions regarding sampling frequency versus sampling duration—bursts of data come at the cost of either sampling duration or frequency (longer gaps in the data). Here we explore how different decisions along this trade-off affect the ability to recover interaction networks. Using three complete days of continuous whole-group GPS tracking of vulturine guineafowl (Supplementary Materials section 6 for methods), we detect all cases of a hypothetical interaction (defined as two individuals within a proximity threshold of 0.6 m for at least 20 s) to construct a ‘true’ social network. We then design a subsampling procedure that collects bursts of data of a given duration (20 seconds to 15 minutes), such that all burst durations approaches collect the same amount of data (25% of the full dataset, or 20 s in every 80 s through to 15 continuous minutes in every hour). Using data from these bursts and applying the same interaction criteria as the true network, we constructed ‘observed’ social networks. This reveals that the capacity to recover a true network (here the correlation between the observed and true networks) varies along the sampling frequency to duration trade-off (Fig. 5), highlighting the importance of design decisions on the resulting study outcomes.

**Figure 5.**
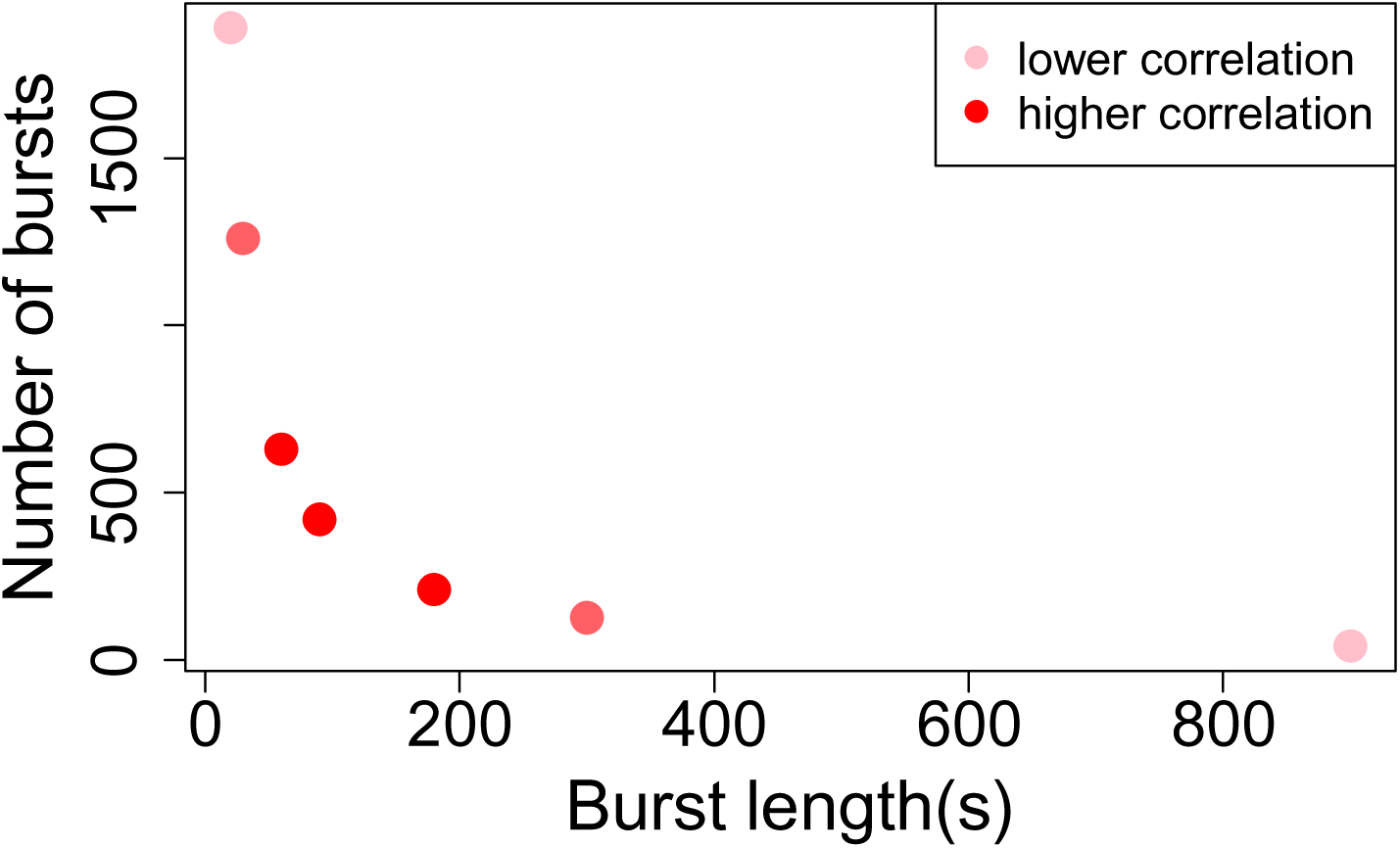
The ability for different sampling regimes—longer bursts versus more frequent bursts— to recover a network estimated from a whole dataset. First, we note that the relationship between sampling duration (here the number of bursts per unit of time, as greater intervals between bursts are often used to extend battery life) and sampling frequency (here burst length) is logarithmic (noting that the precise shape of this relationship will be determined by battery savings arising from collecting data in bursts). Second, the correlation between the observed and the true network varies across this trade-off, and in the case of our hypothetical interaction it peaks at intermediate values for both. While differences in correlation strength in this example were relatively minor (2.3% difference between lowest and highest correlations in our illustrative case study based on only 3 days of data), the pattern is nonetheless robust, and should be expected to produce greater differences in larger datasets. From this analysis, the optimal value yielding the most accurate results was at burst lengths that were 3-6 times longer than the minimum time required for an interaction to take place (20 s).

### 3.3 Constructing social networks

Whereas association events or interactions can be inferred from singular snapshots of spatial positions, social relationships between individuals are typically quantified from the tendency for individuals to associate or interact over (observation) time. The simple ratio index (SRI) defined as the number of observations of two individuals together divided by the total number of observations involving either of them (Hoppitt & Farine 2018), is commonly used to calculate relationship strength. However, this SRI formula needs to be modified for GPS data, as an observation of individual A without individual B is only informative if B is confirmed elsewhere; the absence of data for B provides no information about whether it, or wasn’t, with A. Therefore, the denominator of the formula should only include observations where GPS data are simultaneously available for both individuals.

## 4. SOCIAL BEHAVIOURS WITHIN GROUPS

The application of GPS tracking is particularly promising for studies of animals that live in stable social groups, where the spatial behaviours of group members are typically shaped by those of others (Farine *et al*. 2017). However, it remains unclear how GPS coverage within a social group (which is often the foremost consideration when designing a study) affects the inferences made about individuals’ spatial position in their group and the spatial properties of groups (Fig. 2). To address this gap, we evaluated how sub-sampling influences inferences of individual and group spatial behaviours (see 4.1 and 5.1, respectively), using GPS tracking and simulated data with various sizes and degrees of spatial dispersion of groups (see Supplemental Materials section 6 for methods).

### 4.1 Estimating individuals’ spatial positions within groups

Individuals’ spatial positions relative to other group members can influence vulnerability to predation (Barta, Flynn & Giraldeau 1997), foraging competition with group-mates (Gall & Manser 2018), or availability to receive information from social partners (Rosenthal *et al*. 2015). When inferring such positions, there is a trade-off between the accuracy of estimates and sampling coverage. Here we use empirical and simulated data to show that increasing the proportion of tracked individuals results in exponential decreases in error of estimates of distance to nearest neighbour and spatial centrality (‘surroundedness’). Both empirical and simulated data suggest that, regardless of group sizes, studies can miss up to half of the individuals in a group while still being able to make accurate estimates of nearest-neighbour distances (i.e., median error <1 m when 50% tracked, Fig. 7-a, i-iii). However, accuracy is also affected by group cohesiveness, with less cohesive groups requiring a significantly larger sampling coverage to capture similar accuracy (Supplementary Materials section 7). While errors are generally small, it is worth noting that any non-zero error would also result in mis-assignment of the identities of individuals’ neighbours (Fig. 6; the chance of mis-assignment is the same as the proportion of non-tracked individuals in groups). For surroundedness (Fig. 7-b, i-iii), reliable estimates require a larger proportion (≥60%) of a group to be GPS-tracked, especially in smaller groups, and spatial dispersion has relatively little effect (Supplementary Materials section 7). The relationship between GPS coverage and error in estimates of nearest neighbour distance and spatial position is exponential (see Table S1), and, in both cases, lower GPS coverage is necessary in larger groups.

**Figure 6.**
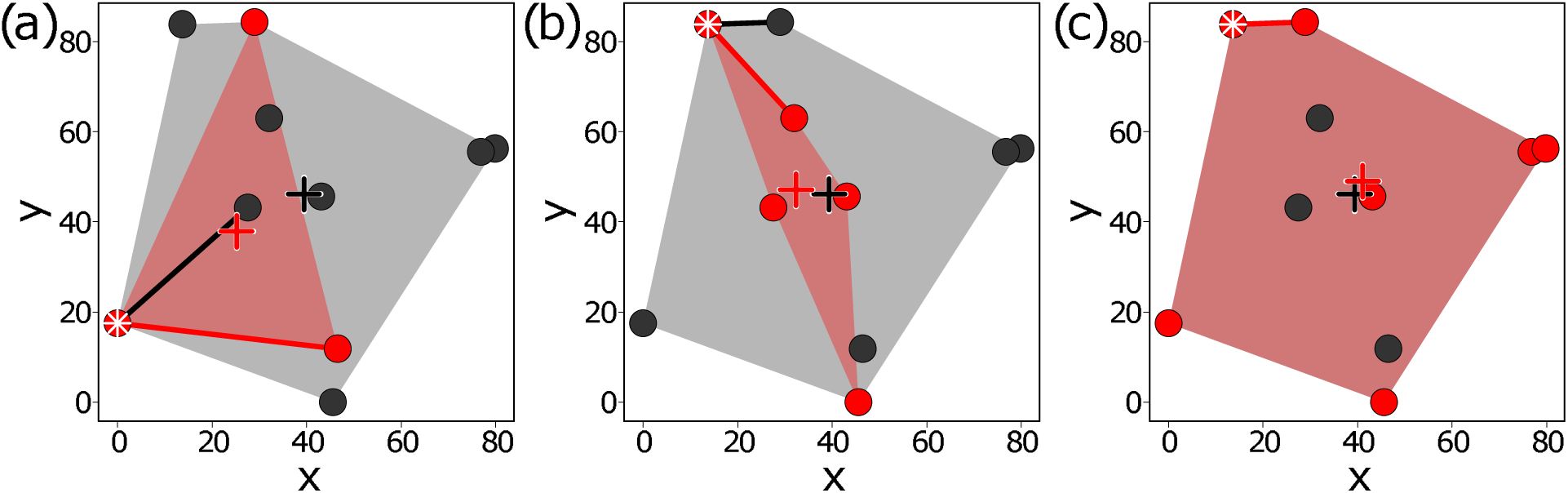
An illustration of the effects sub-sampling on estimates of individual- and group-level properties. Deploying GPS devices on a subset of group members (black: un-sampled group members, red: sampled individuals) can impact behavioural measures. At the individual-level, subsampling influences the accuracy of estimates in the identity of, and distance to, the nearest neighbours (e.g., red focal individuals marked with white stars in a–c in subsamples have different nearest neighbours and larger nearest neighbour distances—segments in black and red are the true and observed patterns, respectively); at the group-level, subsampling influences the accuracy of estimates in group spatial coverage (i.e., MCP, the minimum convex polygons, a–c) and the geometric centroid of the spatial positions of individuals (crosses, a–c).

**Figure 7.**
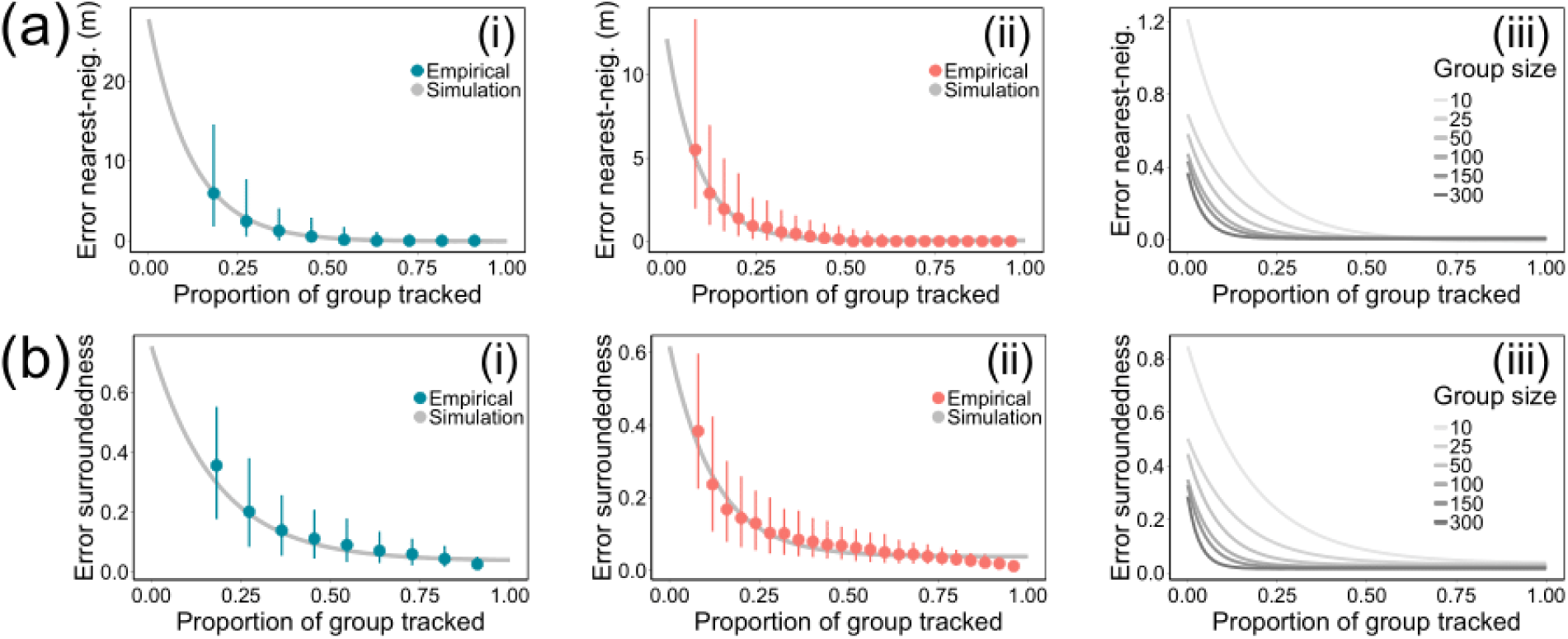
Error in estimates of the nearest-neighbour distance (a) and surroundness (b) decrease exponentially with increasing GPS coverage. (i-ii) Tracking data from two vulturine guineafowl groups (i: N = 11, blue; ii: N = 25, red) randomly subsampled to explore the effects of GPS coverage on estimates of individuals’ spatial positions in groups. Points represent the median error of estimates from the subsampled empirical data, with vertical bars showing the 1^st^ and 3^rd^ quartiles of errors. Curves represent the predicted median error of estimates from exponential models (Table S1) fitted to simulated data matching the spatial extents and group sizes in empirical data. (iii) Curves showing the exponential relationship between GPS coverage and error drawn from simulated data with varying group sizes (Table S2).

### 4.2 Collective movement and decision-making

A typical characteristic of group-living animals is the emergence of collective behaviours (Sumpter 2006). For instance, groups of fish engage in collective schooling behaviour to reduce predation risk to any individual (Herbert-Read *et al*. 2017), while griffon vultures (*Gyps fulvus*) forage in a ‘sky network’ where individuals learn about available food sources by observing the behaviour of their neighbours (Cortés-Avizanda *et al*. 2014). Studying collective behaviours requires perhaps the greatest observational effort (Strandburg-Peshkin *et al*. 2015; Hughey *et al*. 2018), comprising high-spatiotemporal-resolution movement data (high sampling frequency) and high GPS coverage in order to capture the transient socio-spatial dynamics that produce collective outcomes. To date, most studies have been conducted in semi-wild systems, such as homing pigeons (Pettit *et al*. 2013; Sankey *et al*. 2021), where individuals can be easily recaptured to apply and remove GPS devices. However, the limitations of sampling frequency can be overcome by focussing sampling effort to discrete periods to maximize the collection of simultaneous high-resolution data. For example, solar-powered GPS devices on vulturine guineafowl are given three days to recharge so that all devices have full batteries to collect high-resolution at the same time every fourth day (Box 1). Further, some consequences of collective behaviour may be able to be studied using a subset of group members. For example, how collective movements are affected by social (e.g., group size, Papageorgiou & Farine 2020a) or environmental conditions (e.g., seasonal rainfall, Papageorgiou *et al*. 2021), and how moving within a group affects individuals’ behaviours, such as by imposing mechanical (Harel, Loftus & Crofoot 2021) or energetic (Klarevas-Irby, Wikelski & Farine 2021) constraints on movements (Klarevas-Irby, Wikelski & Farine 2021), could be estimated from the tracks of single individuals when groups move cohesively.

## 5. BEHAVIOURS AMONG GROUPS

Assessing behaviours at the group level can reveal inter-group behavioural variability and the influences that groups exert on each other. One prominent way in which groups vary is in their movement characteristics—for instance, meerkat groups emerge from their burrows at different times of day (Thornton, Samson & Clutton-Brock 2010), and vulturine guineafowl groups differ in home range size and daily travel distance (Papageorgiou & Farine 2020a). At its simplest, in cohesive groups with stable membership, the group’s movement can be estimated from a single GPS-tracked group member (Papageorgiou & Farine 2020a; Papageorgiou *et al*. 2021). This can, for example, capture responses to inter-group encounters (Crofoot *et al*. 2008; Papageorgiou *et al*. 2019) or to dispersing individuals (Alberts & Altmann 1995; Armansin *et al*. 2020; Klarevas-Irby, Wikelski & Farine 2021). However, there is currently little guidance on how many GPS-tracked individuals are required to accurately characterise some key group-level properties. Metrics such as the precise location of the group (e.g., for habitat selection of groups) or the level of spatial dispersion of group members (e.g., as an indirect measure of risks) will be more precise when more individuals are tracked. Further, studying behaviours such as inter-group interactions or associations—which are substantially rarer than within-group interactions (e.g., when only a small proportion of individuals in each group infrequently interact or associate with others from different groups)—will generally require deploying more GPS devices per group (Box 1) and increasing GPS sampling duration. Here we consider some of these measures for groups that maintain stable and cohesive membership.

### 5.1 Behavioural characteristics of groups

Group location (i.e., the geometric centroid of all individuals) and spatial dispersion (i.e., the total area covered by a group) are fundamental metrics characterizing group-level behavioural patterns. Using both empirical and simulated data (Supplemental materials section 6), we find that estimating the precise location of cohesive groups is relatively robust to low GPS coverage. In vulturine guineafowl, as few as two tagged individuals can estimate a group’s centroid with <5 m of error (Fig. 8a, i-ii), which is small given that groups can travel several kilometres per day (Papageorgiou *et al*. 2021). For both simulated and empirical data, the relationship between GPS coverage and centroid estimation error follows a negative exponential relationship (Fig. 8a). As such, unless high precision is needed, the spatial location of groups can be sufficiently estimated with as few as 1-3 devices per group, regardless of group size. This is partly affected by group dispersion, with spatially more dispersed groups requiring larger sampling coverage than spatially more cohesive groups to achieve the same precision (Supplemental Materials section 7). Further, we note that in more dispersed groups, the group centroid is less likely to capture where any individual is likely to be.

**Figure 8.**
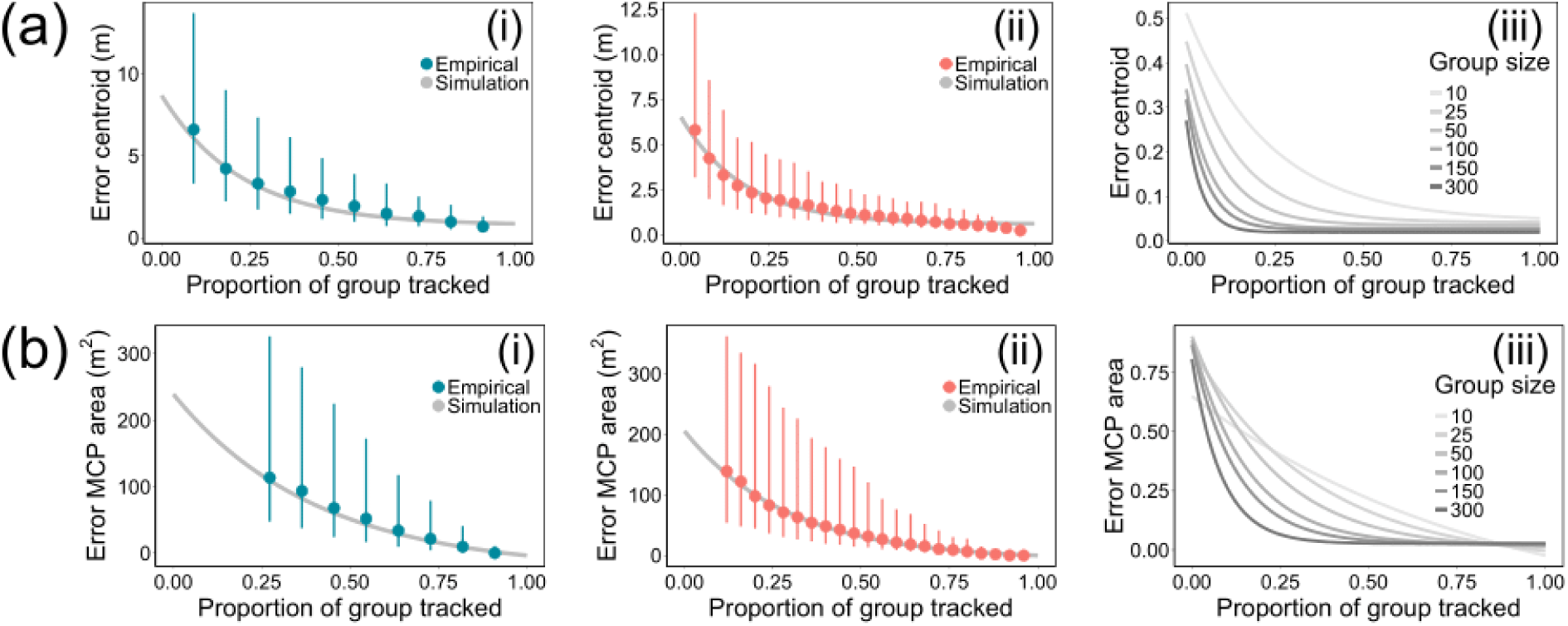
Error in estimates of group centroid (a) and spatial dispersion (b) in cohesive social groups is affected by the proportion of individuals that are tracked. Panels (i) and (ii) show estimates from two vulturine guineafowl groups (i: N = 11, blue; ii: N = 25, red) and simulated data randomly subsampled to explore the effects of GPS coverage on estimates of group position (i.e., centroid, a) and dispersion (i.e., the area of a MCP, b). Points represent the median error of estimates from the subsampled empirical data, with vertical bars showing the 1^st^ and 3^rd^ quartiles of errors. Curves represent the predicted median error of estimates from exponential models (Table S1) fitted to simulated data matching the spatial extents and group sizes in empirical data. Panels (iii) showing the exponential relationship between GPS coverage and error drawn from simulated data with varying group sizes (Table S2).

Estimating a group’s spatial dispersion [e.g., the area of a minimum convex polygon (MCP) containing all group members] requires the majority of the group to be marked, especially in smaller groups (Fig. 8b, i-iii). Even with very high GPS coverage (e.g., >80%), measures often underestimated the dispersion of the group by 20% or more. The less cohesive (i.e. more dispersed) a group, the more sampling coverage is needed to achieve the same accuracy (Supplemental Materials section 7). The relationship between GPS coverage and spatial dispersion error is well captured by a negative exponential function (Fig. 8b).

### 5.2 Inter-group relationships

Groups influence each other through the dispersal of individuals (Alberts & Altmann 1995; Bond *et al*. 2021), associations between groups (Qi *et al*. 2014; Papageorgiou *et al*. 2019; Henriquez *et al*. 2021), or agonistic interactions (Thompson *et al*. 2017; Dyble *et al*. 2019). Studying dispersal requires deploying GPS devices to both dispersers and to the groups that these dispersers might move between. In such cases, a single tagged resident individual in each group is likely to be sufficiently informative if group membership is highly stable. Studying intergroup associations, where membership is stable and groups are cohesive, follows the same principles as studying associations among individuals (see 3.1), where the unit of focus (i.e., groups) can be meaningfully represented by a single individual (if the scale over which groups are spread is small relative to the space over which groups move). Finally, while studying agonistic interactions among groups broadly follows the same principles as studying interactions among individuals, there may be additional challenges. Groups may detect each other over large distances and be more likely to passively avoid interactions (e.g., through scent marking; Christensen *et al*. 2016), leaving the spatial scale at which groups interact unclear. Further, not all individuals might contribute equally to inter-group interactions (Gavrilets 2015), meaning that consideration should be put regarding who to deploy GPS devices onto.

## 6. SOME STRATEGIC CONSIDERATIONS

Besides the constraints outlined above, several other factors should be considered when deploying GPS devices to study social behaviour. Below we list and discuss a few of these, and in Box 2 we outline how we deployed GPS devices and programmed these to achieve the multiple aims of our vulturine guineafowl project.

### 6.1 Redundancy

In most cases, studies should consider tracking extra individuals in each group beyond the minimum required for the study design. For example, while the locations of each cohesive social group might be captured by a single group member, its GPS device could fail, fall off, or the individual could die. Further, when using mixed GPS sampling strategies (see 6.3), having multiple individuals in each group can maximise the probability of at least one individual collecting higher resolution data.

### 6.2 Partitioning sampling effort in time

It can be useful to vary GPS sampling across time to focus data collection on certain time windows over others. For example, diurnal species are largely inactive at night (Isbell *et al*. 2017; but see Loftus *et al*. 2021), meaning that GPS devices are often switched off at night to conserve battery. It can also be worthwhile to program devices to increase sampling frequency during periods when animals are more active (e.g., morning and/or evening), or decrease sampling frequency during seasons when conditions are less optimal for data collection (e.g., during breeding seasons, Box 2). Some tags can activate higher resolution sampling using sensors that require less energy (e.g. accelerometers). However, when partitioning data collection, it is worth considering whether partitioning sampling in time could introduce biases or result in incorrect inferences (see, for example, section 3.1).

### 6.3 Mixed GPS sampling frequencies

The major trade-off is between sampling frequency and sampling duration. Even with solar-powered GPS devices, the capacity to collect high-resolution data across entire days is limited. In stable, cohesive groups, a solution to this is to use a mixture of sampling regimes—having some individuals deployed to prioritise shorter bursts of higher sampling frequencies and some individuals prioritised to capture long-term sampling. With solar-powered devices, it is often possible to programme devices to switch from low– to high-resolution as the battery charges (Box 1). Such an approach can allow devices deployed across a group to work in a collaborative way.

### 6.4 Supplementing GPS tracking with other observational data

A key limitation of GPS tracking data for inferring social behaviours is that researchers are blind to the absence/presence of untracked individuals in close spatial proximity. GPS tracking studies can therefore benefit from supplementary observational data (Smith & Pinter-Wollman 2021). Behavioural patterns revealed by observational data can be the ground-truth to those inferred with GPS tracking data and can be used to evaluate the extent to which data derived from a given deployment of GPS sampling are able to capture true behavioural patterns in animal societies. However, integrating GPS and non-GPS data into the same analysis introduces data compatibility challenges, and approaches for harmonizing such datasets warrant further development.

### 6.5 Combining GPS data with other sensors

Modern GPS tracking studies use additional sensors (Williams *et al*. 2020), such as accelerometers, gyrometers (Marin 2020), heart rate monitors, or body temperature loggers (e.g., Linek *et al*. 2021). Such data can substantially increase the dimensions of information researchers can retrieve about animals’ behaviours (e.g., Harel, Loftus & Crofoot 2021). For example, movement-related sensors can improve data on trajectories by allowing tracks between GPS fixes to be reconstructed using dead reckoning (Wensveen, Thomas & Miller 2015), while the fluctuations in on-board battery voltage of solar-powered GPS devices could potentially be used to infer environmental conditions social animals experience (e.g., whether an animal is under cover or not).

In social contexts, accompanying data from accelerometers can also be useful when inferring dyadic social interactions such as inter-individual displacements (Strandburg-Peshkin *et al*. 2015), individual behavioural states (Chimienti *et al*. 2016), or behavioural synchrony among individuals (Strauss *et al*. 2021). For example, accelerometers can capture if two individuals are stationary or not, thereby allowing their estimated distance to be more accurately calculated by averaging their positions over consecutive fixes. Acceleration data could also potentially be used to infer social behaviours, but a key challenge is the interpretation or classification of those behaviours, which encompass large locomotor and behavioural diversity (Shamoun-Baranes *et al*. 2012; Brown *et al*. 2013). Automated analytical tools such as machine learning have been developed to identify behavioural states (Fehlmann *et al*. 2017; Chakravarty *et al*. 2019) and could in theory be applied to social behaviours, especially, if they play out over longer periods (e.g., allo-grooming or -preening) and are characterised by some stereo-typed body movements. One challenge, however, is that many social interactions are brief (e.g., displacements, greeting behaviours), executed in a variety of ways and can resemble non-social behaviours (e.g. running from conspecifics versus from predators), thus resulting in small or noisy datasets. The social context of pure accelerometer-derived behaviours can also be ambiguous, for example, two individuals detected as moving simultaneously could be doing so together or apart. Thus, high-resolution GPS data might, in turn, be important for inferring whether behaviours are being expressed socially or not. More work is needed on developing accelerometer-based methods to help with studies of social behaviours.

The potential benefits of additional sensors also need to be weighed against some costs. For example, additional sensors can increase battery consumption (albeit often a lot less than the GPS sensor itself), and can substantially increase the cost of data storage, retrieval, and management, especially in studies generating large volumes of data (Box 1). Furthermore, although some devices come with multiple sensors onboard (e.g., combined GPS sensor and accelerometer), others might require integrating a separate device along with the GPS device. This can be physically challenging (e.g., in small animals there may not be room for a second sensor or device, or exceed weight limits), and for studies requiring synchronization between data streams, communication must be established between the devices or sensors.

## 7. CONCLUDING REMARKS

GPS tracking is a powerful tool for studying social behaviour, but sampling design involves many trade-offs. Drawing from previous studies, and empirical and simulated data, we have discussed how sampling coverage, sampling duration, and sampling frequency can be optimised to address different research questions. For example, many social measures—of individuals within groups (e.g., spatial position) or of groups themselves (e.g., centroids)—can be reliably estimated by tracking relatively few group members. Other measures, such as social interactions among individuals, social processes, or the spatial dispersion of groups, require more intensive sampling frequencies or greater coverage of individuals. Further, estimations involving calculations of inter-individual distances need to consider the persistent overestimation bias introduced by error in recorded GPS positions.

In terms of overcoming the sampling frequency-duration trade-off, we caution against aiming to sample at the highest frequency possible at the cost of sampling duration. We demonstrated how balancing these can yield more accurate results when studying interactions. A good rule of thumb may be to sample at a frequency that is slightly longer than the behaviour under study (Fig. 5), though this warrants a more focused study addressing the generality of this finding. For other metrics, such as associations, the design should consider at what rate social behaviours might change over time.

Although our discussion here has focused on groups with stable group membership, some of our findings may also apply in more open groups. However, open societies with fluctuating group memberships and sizes present a different challenge for GPS-based research, as the pool of individuals from which to sample is often unpredictable. Our findings at the group level are likely to be relevant when considered at the scale of whole populations, and the framework we apply in this paper to examine the robustness of GPS-based inference about social groups could be used to test, and guide, the design of similar studies in open animal societies.

### Box 1.

The GPS-based vulturine guineafowl project was established at Mpala Research Centre in central Kenya in 2016 to study social dynamics within and across groups, at timescales spanning from moment-by-moment movement to lifetime ranging. A pilot study on group membership of a small number of colour-banded individuals and five GPS-tracked birds (15 g, solar-powered devices, e-obs Digital Telemetry, Grünwald, Germany) showed that group compositions and sizes were highly consistent. The pilot study also confirmed the advantages of using solar-powered GPS devices in an equatorial savannah environment—solar conditions there allow for consistent GPS sampling throughout the year from 06:00 to 19:00. Subsequently, more solar-powered GPS devices were deployed to individuals both within and across groups. Specifically, we aimed to track one bird for every ten group members, allowing us to distribute sampling effort across as many groups as possible (eventually with a coverage of an entire population consisting of >20 groups). GPS sampling effort was designed to prioritise simultaneous population-level behavioural patterns (see section 3.1), by programming the devices to switch on for a 10-second sampling window every 5 minutes (i.e., the ‘Population regime’, Table 2). To take advantage of solar power, devices were programmed to sample at high frequency (i.e., 1 Hz sampling for 15 minutes at a time) when battery charge was high. Every bird in each group was also colour-banded to facilitate visual identification in the field, regardless of whether they were GPS tagged or not. Combining twice-weekly field observations of group membership and size with our population-level GPS data shed light on the underlying patterns of social associations among individuals (both within and across groups), revealing the multilevel social structure of the whole population (Papageorgiou *et al*. 2019). Over time, continued measurements further revealed how group size affects movement and ranging (Papageorgiou & Farine, 2020a) and how groups use the landscape across seasons (Papageorgiou *et al*. 2021).

**Table 2.**
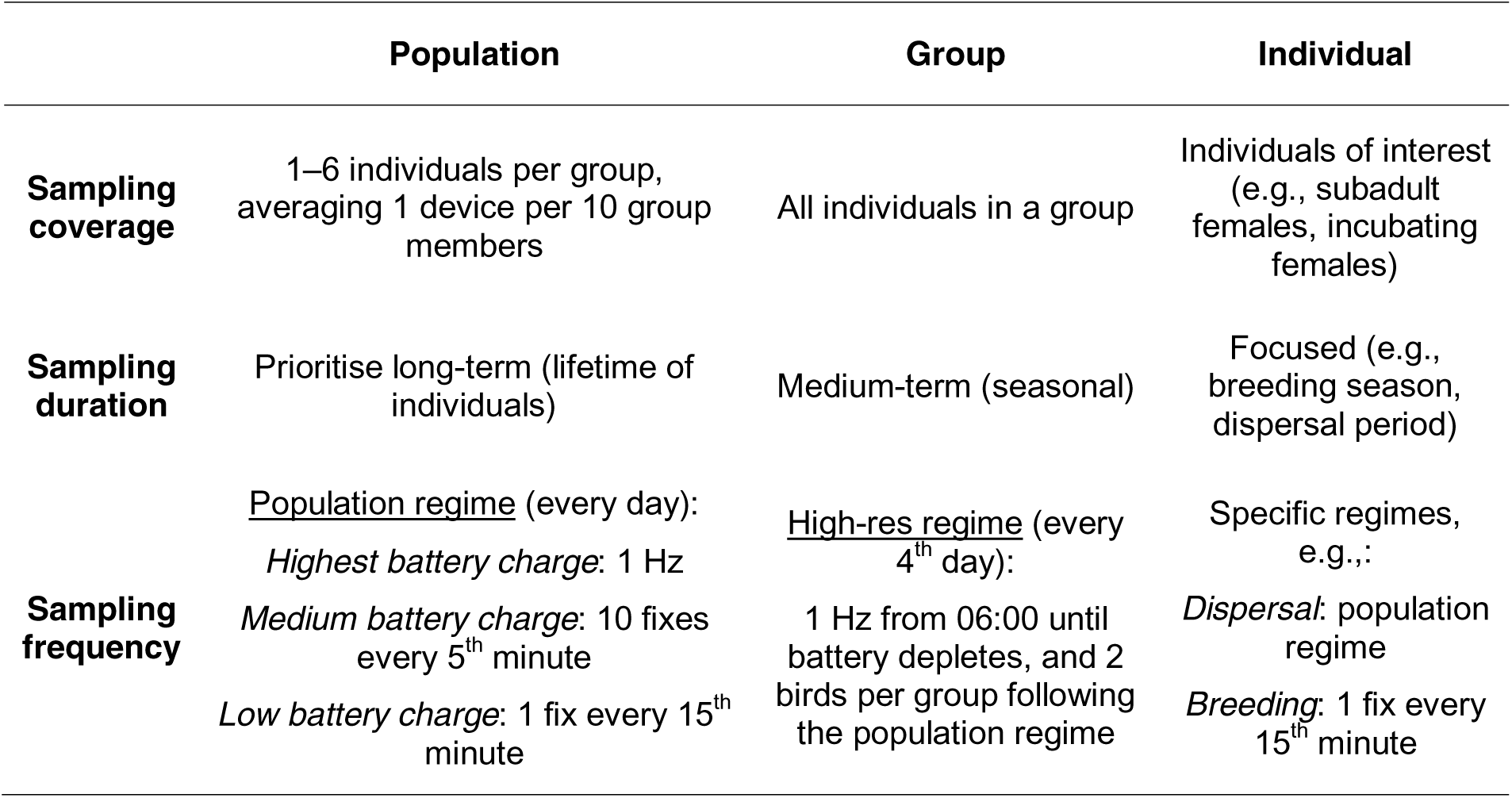
Overview of GPS sampling strategies for the vulturine guineafowl project.

Under the population-level sampling regime (Table 2), devices collect data at high sampling frequency approximately 25% of the time, suggesting that it is feasible to collect extensive movement data with high temporal resolution. However, variation in battery charge can lead to asynchrony in high-resolution data collection across devices. To study within-group social dynamics (e.g., how groups make decisions), every bird in several focal groups were GPS-tracked and devices were programmed to collect data on the same day, every fourth day, spending the intervening days recharging (except for one or two birds in each group with devices programmed with the population regime to track daily ranging). This ‘high-res’ regime (Table 2) ensures that all devices collect >6 h of synchronous, fine-temporal-scale movement data of all individuals on each operating day (albeit potentially biasing observations to morning rather than evening behaviours). Such data have supplemented observational data to infer how individuals respond to group decisions (Papageorgiou & Farine 2020b), and to characterise other aspects of group behaviour, as in this study.

Field observations suggest that males are philopatric while females disperse (Klarevas-Irby, Wikelski & Farine 2021). Thus, devices were primarily deployed onto males for long-term data collection. As some adult females bred, GPS devices were also deployed to these females for opportunities to find nests and to study incubation behaviours. However, the batteries of devices deployed on these breeding females often struggled to maintain an adequate charge due to poor solar conditions— incubating birds can stay under dense vegetation for up to 25 days at a time (Nyaguthii 2021). We benefited from the ability to remotely program devices—reducing sampling frequency to one fix per 15 minutes ensured the capacity of batteries to cover the whole incubation period. To study dispersal behaviours, subadult females were GPS-tracked (Klarevas-Irby, Wikelski & Farine 2021). In this case, new devices were deployed to ensure maximum battery power (sampling at high frequency covering ca. 30% of the time in a day). Comparing fine-scale movements of dispersers with residents relied heavily on the ability to have at least one device in each group sampling at high frequency at any time (facilitated by having multiple tagged individuals per group).

Overall, the project has had as many as 129 simultaneously GPS-tracked birds, and many challenges have arisen in project maintenance and data management. For example, each device produces ca. 4 MB data that scales up to >500 MB data every 10^th^ day across the whole population, requiring up to 4 h of downloading time every second day. This is in addition to extensive effort and time needed to regularly check and maintain deployed devices, and to recover devices when individuals are predated. To partially address these challenges, the accelerometer sensors integrated into our devices have been turned off, as the resulting data streams would prove infeasibly large for our current download regime (but the devices can be re-programmed for collecting acceleration data if desired).

## ACKNOWLEDGEMENTS

We thank the e-obs GmbH for technical advice on GPS battery life management, the vulturine guineafowl field team coordinated by Brendah Nyaguthii in Kenya, and two anonymous reviewers for their constructive comments on the manuscript. Data were collected with permission from, and in collaboration with, the Kenya National Science and Technology Council (NACOSTI/P/16/3706/6465), the National Environment Management Authority (NEMA Access Permit NEMA/AGR/68/2017), the Kenya Wildlife Service (KWS-0016-01-21), the Wildlife Research Training Institute (WRTI-0026-02-21), the National Museums of Kenya, Dr. Peter Njoroge, the Mpala Research Center. Ethical approval was granted by the Max Planck Society’s Ethikrat Committee. This study was funded by a grant from the European Research Council (ERC) under the European Union’s Horizon 2020 research and innovation programme (grant agreement No. 850859), an Eccellenza Professorship Grant of the Swiss National Science Foundation (Grant Number PCEFP3_187058) awarded to DRF, and the Max Planck Society. PH, JAKI, and DP acknowledge support from International Max Planck Research School (IMPRS) for Organismal Biology. EDS was funded by the Alexander von Humboldt foundation.

## AUTHORS’ CONTRIBUTIONS

PH, JAKI and DRF conceptualized the study. CC collected data for testing GPS precision. PH, JAKI and DRF conducted the analyses. PH drafted the manuscript and led the revisions. All authors contributed critically to the revisions and gave approval for publication.

