## Supplementary materials for "A guide to sampling design for GPS-based studies of animal societies"

### 1. Practical considerations when studying animal societies using GPS tracking

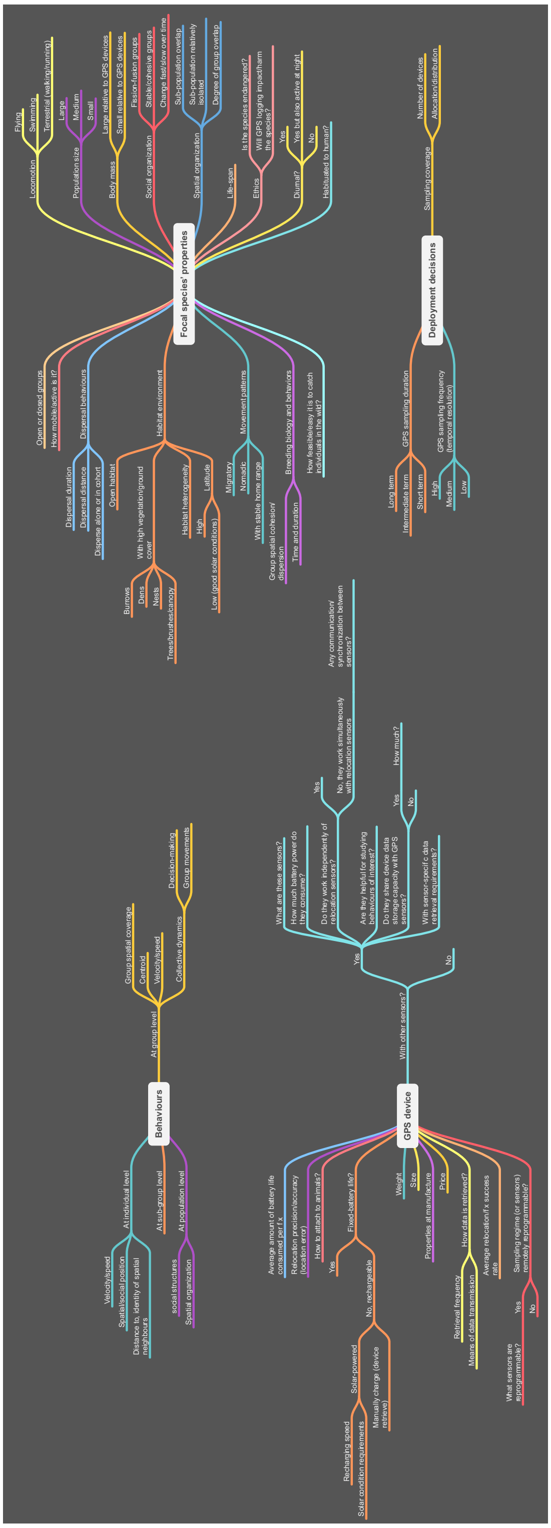


**Figure S1. GPS deployment considerations in relation to behaviours of interest, the biology or social system of focal species, and the properties of GPS devices, etc.**

#### 2. Empirical and on-board estimations of GPS error

GPS tags often provide an on-board estimation of error. To test how informative this measure is, we compared it to the GPS error obtained from the field test of the 15g e-obs solar tags, where tags set to record at 5-, 10-, 30- and 60-minute intervals recorded data for 8 consecutive days. While the error estimation obtained from field testing was comparable to the on-board GPS accuracy, the absolute values differed substantially (Fig. S2).

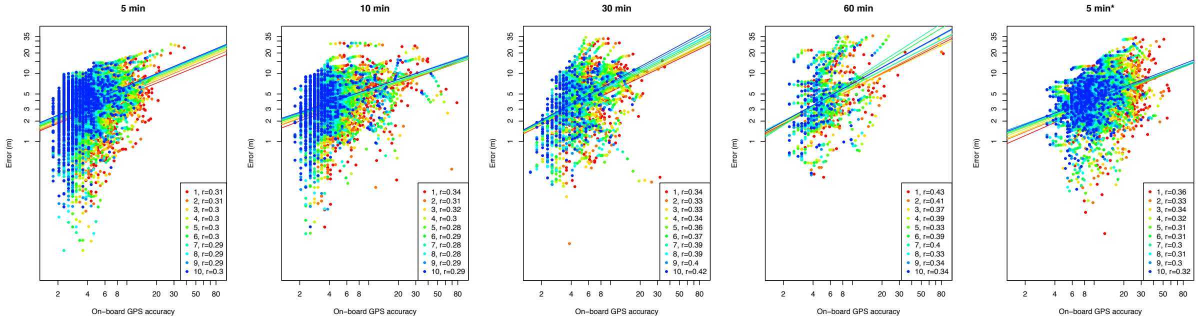


**Figure S2. On-board estimations of error associated with each fix holds minimal information.** Plots are shown for different sampling regimes (and for canopy cover marked by a "*"). These show that while the error estimated by the tag correlates with the on-board GPS estimation of positional error for that fix, the correlation is low, making the on-board information mostly uninformative (note the log-log scale).

#### 3. Theoretical predictions for dyadic distances

To demonstrate the effect of GPS accuracy (i.e. errors in the estimated position of each fix), we generated two paired sets of X, Y coordinates with their mean at a set distance but with error (standard deviation of a Gaussian distribution). We then calculated the proportion of pairwise detections that were within the true distance as a function of the position of the observe points (Fig. S3-a). Repeating this process across a range of true distances and tag errors shows that the estimated mean distance will be consistently larger than the true distance, but vary according to tag error and true distance (Fig. S3-b).

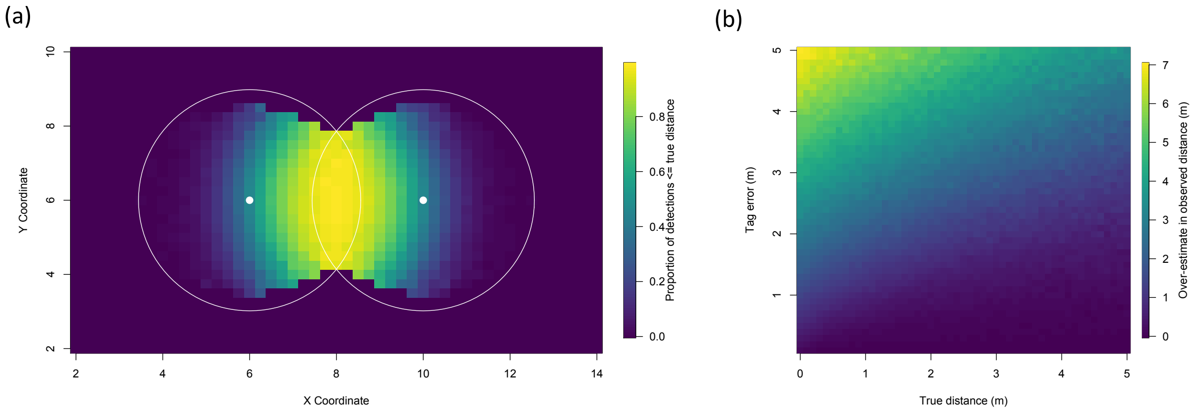


**Figure S3. Relationship between tag error and estimation of inter-individual distances.** (a) Two individuals that are positioned a fixed distance apart (represented by two white dots) with tag error (distributing points according to the white circles) will have a greater proportion of their simultaneous detections separated by a greater distance than their true distance, as demonstrated by the bias in positive detections (within or equal to the true distance) in the direction of the other individual. (b) The resulting over-estimation in inter-individual distance increases as a function of tag error, but decreases as a function of the true distance between individuals.

#### 4. Removing GPS precision confirms the predicted relationship between the over-estimate in inter-tag distance and the true distance

To confirm that the deviation in the relationship between the inter-tag distance and true distance in the empirical data is due to GPS precision (the correlation in errors between tags), we randomised the timestamps in the data used to create Fig. 4. This effectively removes GPS precision (between-tag similarity in errors, or error in relative space), but keeps GPS accuracy (the error for each unique fix in true space). This analysis confirms the theoretical relationship, and therefore that within-tag inaccuracy may not have as much of an impact on estimates of inter-tag distances, especially at closer distances.

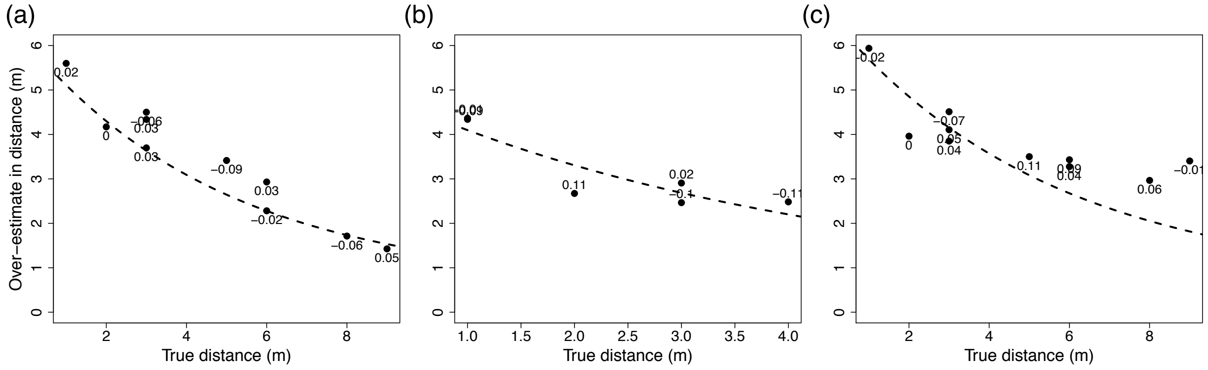


**Figure S4. Relationship between over-estimate in inter-tag distance and true inter-tag distance when removing GPS precision.** (a) GPS-based distance calculations consistently over-estimated the true tag distance, and when GPS precision is removed the relationship matches the theoretical prediction (dashed line). (b) Height above ground and tag being 'on the move' reduces the GPS accuracy error also matches the theoretical prediction when removing GPS precision. (c) Canopy causes a constant over-estimation that deviate from the predicted relationship at larger true distances.

#### 5. Estimating true association rates

Imagine that a researcher is interested in capturing the proportion of time that two individuals are within a given meaningful distance (e.g. to detect physical contact or grooming interactions, which we call a biological threshold). As a consequence of the over-estimation in distance (Fig. S5), there will be an increase rate of false negatives (true distance is within the biological threshold but the detection is not) as the true distance approaches the biological threshold (Fig. S5-a). This would persistently under-estimate the proportion of time individuals spend within the biological threshold, i.e. their association strengths. Because the relationship between true and inferred inter-individual distance is relatively predictable, it may be meaningful to increase the acceptance threshold for associations. Doing so accepts some false positives (true distance greater than the biological threshold but the detection is not), but can do so at a rate that matches the false negatives (Fig. S5-b), thereby balancing these rates so that across true distances, the association rate stays constant. This alternative threshold would vary according to both the desired biological threshold and the tag error (Fig. S5-c).

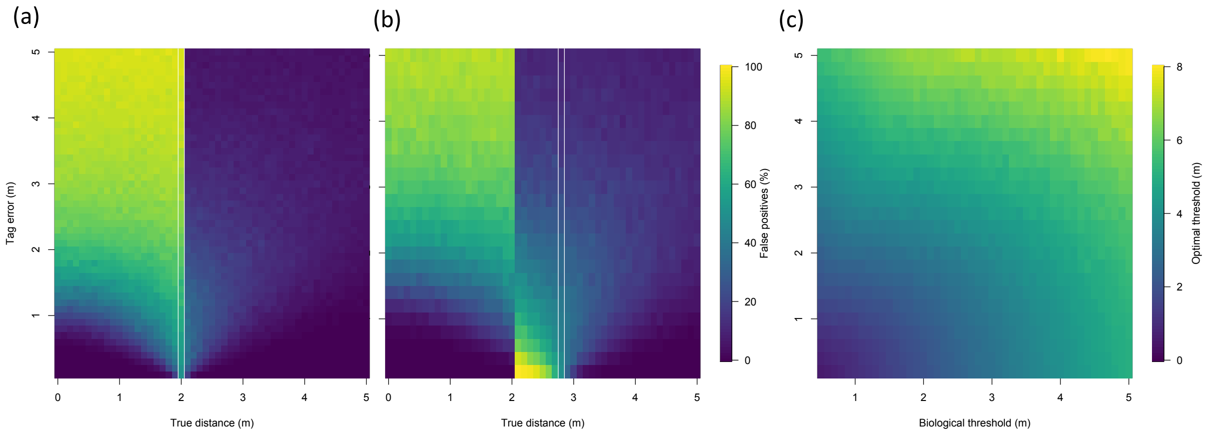


**Figure S5. Estimating social connections from GPS-based inter-individual distances is prone to error, and the detection threshold required for accurate estimates of association rates may be larger than the distance estimated for biologically meaningful detections.** (a) The rate of false negatives (X ≤ 2) and false positives (right of X > 2) as a function of the true inter-individual distance and tag error, when a detection threshold is set at exactly the biological threshold (here 2 m). (b) Corresponding error rates (false negative when X ≤ 2 and false positive and false positive when X > 2) when the detection threshold is set above the biological threshold (here 2.8 m). (c) Example of an adjusted detection thresholds for each given combination of biological thresholds and tag errors, illustrated for an optimal threshold is defined as the detection threshold where the probability of a false negative is equal to the probability of a false positive at or just above the biological threshold, respectively.

#### 6. Supplementary methods for empirical data collection and simulations

Our empirical data is drawn from two social groups of vulturine guineafowl (N = 11 and N = 25) in which all members were simultaneously tracked. Each bird in each group was fit with an e-obs 15g solar GPS tag set to collect one point every second (1Hz) on every fourth day (see Box 1 in the main texts). For the analyses in section 3.3, we selected the first three days of whole-group GPS tracking, which we subsampled as described in the main text.

For the analyses in sections 4 and 5, we randomly selected 100 time stamps from the tracking data, spanning several months, from each group. At each time stamp, the data were subsampled by randomly removing group members [producing sub-groups of sizes 1 to (N-1)], with each level of sub-sampling repeated 10 times (i.e., producing 10 combinations of subsampled individuals for each time stamp). We then extracted four spatial measures describing individual positions and group properties: the distance from some focal individual to its nearest neighbour (a common measure of social association), the spatial centrality of a focal individual within the group (i.e., 'surroundedness', or 1 – CV, where CV is the circular variance of the vectors pointing to the focal individual; for CV see Christman & Lewis 2005 in *Animal Behaviour* 70-1, pp. 73-82), the spatial position of the entire group (the geometric centroid of all sampled individuals' positions), and the spatial spread of the group [represented by the area of the minimal convex polygon (MCP) containing all sampled individuals]. The effect of sub-sampling on each measure was quantified as the difference between the "true" value derived from the full group data and the "observed" value derived from the subsampled individuals only. For nearest-neighbour distance and centroid, the error was defined as the point-to-point distance from the true position (of the spatially-nearest neighbour or the original centroid, respectively) to the observed position (of the nearest remaining neighbour or the new centroid). Error in surroundedness was defined as the difference between the true and observed circular variance for some focal individual included in the subsampled group. Error in group spread was defined as the difference in the areas of the MPCs between the subsampled and original group.

We then simulated data to confirm the relationships between GPS coverage and error. First, for each time stamp in the empirical data, we simulated empirically derived groups (N = 11 and N = 25), where simulated individuals' X and Y coordinates were drawn from uniform distributions nested within the rectangles outlined by each empirical snapshot. We simulated 100 empirically derived group observations to match each time stamp in the empirical data, and subsampled and measured these following the same procedure as empirical groups.

To examine the general effects of GPS coverage on the accuracy of individual- and group-level behavioural estimates, we complemented our empirical dataset and empirically-informed simulations with purely simulated groups varying in size from 10 to 300. Here, X and Y coordinates were drawn from uniform distributions between 0 and 1. We simulated 100 group observations for each group size, which were subsampled and measured following the same procedure as empirical groups.

#### 7. Effects of group spatial dispersion on estimates of individual- and group-level behaviours

We examined the effects of group spatial dispersion on the accuracy of the estimates of both the individual-level and group-level behaviours. We simulated the spatial positions (i.e., X and Y coordinates) of individuals in three groups (N = 10, N = 50 and N = 100) with varying levels of spatial dispersion—which is characterized by the standard deviations (sd) of the normal distributions of the coordinates nested within the interval [-0.5, 0.5] with 0 as their mean (i.e., a larger sd characterizes a spatially less cohesive group, and vice versa). From the whole dataset, we then subsampled individuals to investigate how tag coverage affects error estimates. In general, we found that the more the individuals in a group are spatially dispersed, the larger the error will be when estimating individual-level and group-level behaviours using a subsample of the group. The exception to this are the estimates of individuals' surroundedness, which do not differ significantly with group spatial dispersion. The effects of group spatial dispersion on the error of behavioural estimates also decrease with group size, and such effects are finite—after reaching a threshold level of spatial dispersion, the error of subsampling tends to converge, no matter how spatially dispersed groups of a given size are.

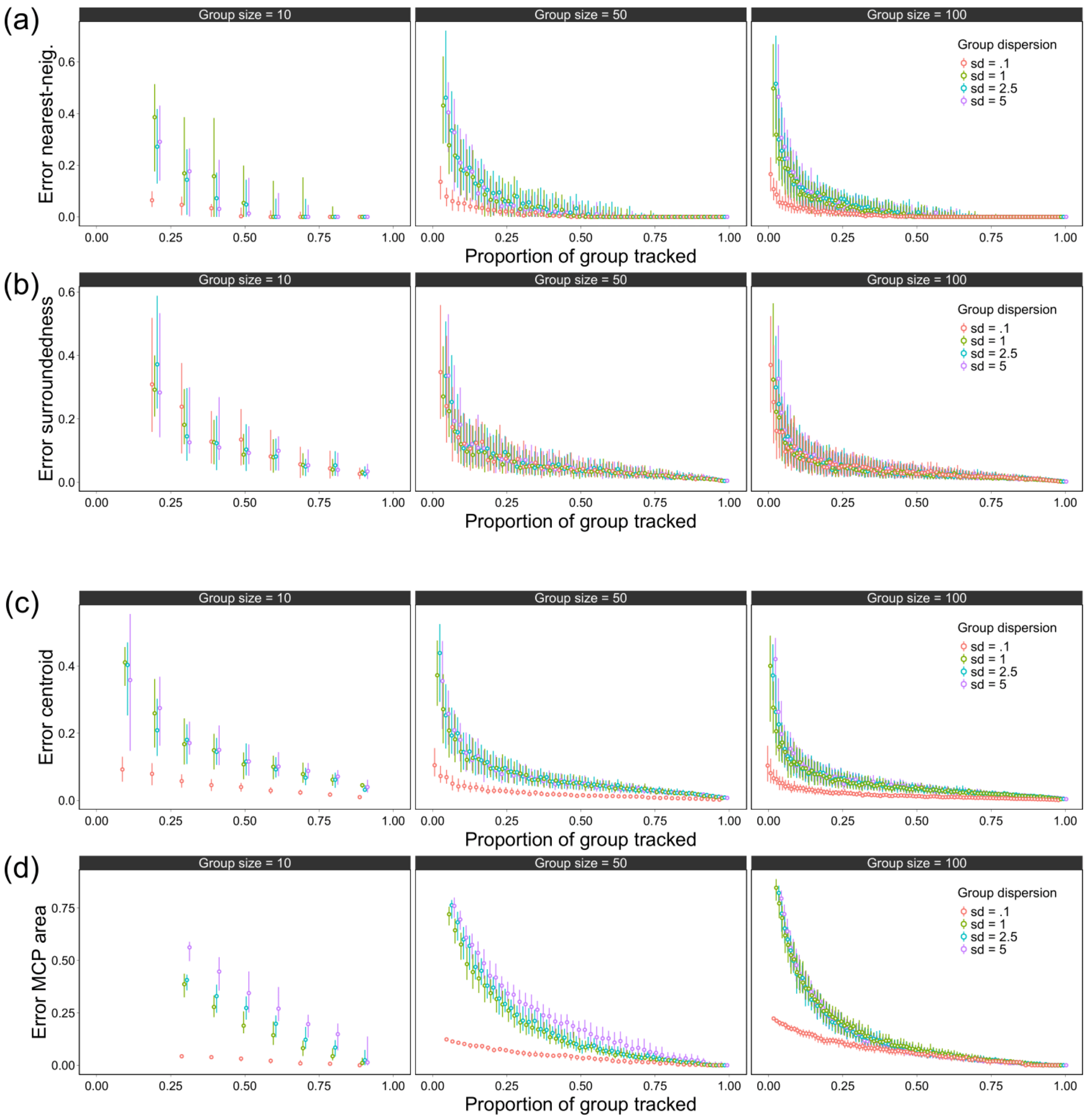


**Figure S6. Effects of group spatial dispersion on estimates of (a, b) individual- and (c, d) group-level behaviours.** Simulations were run for three group sizes (N = 10, N = 50 and N = 100) and varying levels of spatial dispersion (colours). Spatial dispersion was characterized by the standard deviations of the normal distributions (between -0.5 and 0.5 with 0 as their mean) from which individuals' spatial positions (X and Y coordinates) were drawn for behavioural estimates. Circles represent the median error of behavioural estimates from 100 replications in the simulations, while vertical bars mark the 1$^{st}$ and 3$^{rd}$ quartiles of errors.

#### 8. Parameter estimates of the exponential model fitted to the empirical and simulated data

**Table S1. Fit of exponential models to empirical data.** The median error values of each of the four behavioural metrics were fitted to the exponential model $f(x) = \alpha e^{\beta x} + \theta$, where $x$ is the proportion of individuals in a given subsample of two simulated groups of size N, whose size and spatial distributions were modelled after two groups of Vulturine guineafowl (N = 11 and N = 25; see simulation and subsampling methods above). Estimated values of $\alpha, \beta, \theta$ from modelling fitting are given for each group size and metric.

| Metrics | Group size | $\alpha$ | $\beta$ | $\theta$ |
|---|---|---|---|---|
| **Error in nearest-neighbour distance** | N=11 | 27.950 | -8.318 | -0.073 |
| | N=25 | 12.117 | -11.232 | 0.042 |
| **Error in surroundedness** | N=11 | 0.718 | -5.568 | 0.038 |
| | N=25 | 0.578 | -8.005 | 0.038 |
| **Error in centroid position** | N=11 | 7.916 | -4.422 | 0.770 |
| | N=25 | 5.925 | -5.611 | 0.591 |
| **Difference in group MCP area** | N=11 | 265.266 | -2.464 | -26.132 |
| | N=25 | 213.049 | -3.415 | -7.152 |

**Table S2. Fit of empirical models to simulated data.** The median error values of each of the four behavioural metrics were fitted to the exponential model $f(x) = \alpha e^{\beta x} + \theta$, where $x$ is the proportion of tracked individuals in a given subsample of a simulated group of size N (i.e., 10, 25, 50, 100, 150, 300; see simulation and subsampling methods above). Estimated values of $\alpha$, $\beta$, $\theta$ from modelling fitting are given for each group size and metric.

| Metrics | Group size | $\boldsymbol{\alpha}$ | $\boldsymbol{\beta}$ | $\boldsymbol{\theta}$ |
|---|---|---|---|---|
| **Error in nearest-neighbour distance** | N=10 | 1.225 | -6.681 | -0.010 |
| | N=25 | 0.692 | -7.756 | -0.002 |
| | N=50 | 0.577 | -10.649 | 0.005 |
| | N=100 | 0.466 | -13.718 | 0.006 |
| | N=150 | 0.425 | -16.507 | 0.007 |
| | N=300 | 0.361 | -23.227 | 0.007 |
| **Error in surroundedness** | N=10 | 0.812 | -5.560 | 0.035 |
| | N=25 | 0.469 | -7.310 | 0.033 |
| | N=50 | 0.415 | -11.297 | 0.029 |
| | N=100 | 0.325 | -15.344 | 0.023 |
| | N=150 | 0.307 | -19.825 | 0.021 |
| | N=300 | 0.270 | -30.403 | 0.016 |
| **Error in centroid position** | N=10 | 0.467 | -4.303 | 0.044 |
| | N=25 | 0.407 | -6.998 | 0.042 |
| | N=50 | 0.362 | -9.753 | 0.035 |
| | N=100 | 0.312 | -13.55 | 0.028 |
| | N=150 | 0.293 | -17.162 | 0.025 |
| | N=300 | 0.251 | -23.252 | 0.019 |
| **Overlap of group MCP** | N=10 | 0.974 | -1.179 | -0.324 |
| | N=25 | 0.908 | -2.831 | -0.057 |
| | N=50 | 0.903 | -4.361 | 0.002 |
| | N=100 | 0.866 | -6.402 | 0.022 |
| | N=150 | 0.844 | -8.070 | 0.025 |
| | N=300 | 0.782 | -11.993 | 0.026 |
